# A chromosome-level genome assembly enables the identification of the follicle stimulating hormone receptor as the master sex determining gene in *Solea senegalensis*

**DOI:** 10.1101/2022.03.02.482245

**Authors:** R. De la Herrán, M. Hermida, J. Rubiolo, J. Gómez-Garrido, F. Cruz, F. Robles, R. Navajas, F. A. Blanco, P. R. Villamayor, D. Torres, P. Sánchez-Quinteiro, D. Ramirez, M. E. Rodríguez, A. Arias-Pérez, I. Cross, N. Duncan, T. Martínez-Peña, A. Riaza, A. Millán, M. Gut, C. Bouza, D. Robledo, L. Rebordinos, T. Alioto, C. Ruíz-Rejón, P. Martínez

## Abstract

Sex determination (SD) mechanisms are exceptionally diverse and show high evolutionary rates in fish. Pleuronectiformes is an emblematic fish group characterized by its adaptation to demersal life and its compact genomes. Here, we present a chromosome-level genome assembly of Senegalese sole, a promising European aquaculture species. We combined long- and short-read sequencing and a highly dense genetic map to obtain a contiguous assembly of 613 Mb (N50 = 29.0 Mb, 99% of the assembly in the n = 21 chromosomes of its karyotype). The correspondence between this new assembly and the Senegalese sole chromosomes was established by fluorescence *in situ* hybridization with BAC probes. Orthology within Pleuronectiformes was assessed by using the chromosome-level genomes of six important commercial flatfishes covering a broad phylogenetic spectrum of the order. A total of 7936 single-gene orthologues, shared by the six species, were used to identify syntenies in Pleuronectiformes and to explore chromosome evolutionary patterns in the order. Whole genome resequencing of six males and six females enabled the identification of 41 fixed allelic variants in the follicle stimulating hormone receptor (*fshr*) gene, homozygous in females and heterozygous in males, consistent with an XX / XY chromosome system. The observed association between *fshr* SNPs and sex was confirmed at the species level in a broad sample, which allowed tuning up a molecular sexing tool. *Fshr* demonstrated differential gene expression between male and female gonads since 86 days post-fertilization, when the gonad was still an undifferentiated primordium, concomitant with the activation of other testis and ovary marker genes, such as *amh* and *cyp19a1a* genes, respectively. Interestingly, the Y-linked *fshr* allele, which included 24 non-synonymous variants, expressed to a higher level than the X-linked allele at all stages. We hypothesize a molecular mechanism hampering the action of the follicle stimulating hormone that would drive the undifferentiated gonad toward testis.

## Introduction

Sex determination (SD) refers to the mechanism controlling the fate of the gonadal primordium at the initial stages of development responsible for the sex of a mature individual. Despite highly conserved and evolutionary mature SD systems have been reported on mammals, birds and *Drosophila*, the increasing data from ectothermic vertebrates show a sharp different picture (Martínez et al., 2014; Guiguen et al., 2019). Fish display highly diverse chromosome SD systems (Cioffi et al., 2017) and several master SD genes have been reported in this group (Martínez et al., 2014). Among them, classical transcription factors such as *dmy* (Matsuda et al., 2002; Wang et al., 2022), *sox3* (Takehana et al., 2014) or *sox2* (Martínez et al., 2021); transforming growth factor ß-related genes such as *gsdf* (Myosho et al., 2012; Herpin et al., 2021) or *amh* (Hattori et al., 2012; Pan et al., 2019) and its receptor *amhr2* (Kamiya et al., 2012; Feron et al., 2020; Nakamoto et al., 2021; Wen et al., 2022); genes related to the steroidogenic pathway such as *bcar1* (Bao et al., 2019) or *hsd17b1* (Koyama et al., 2019); and finally, some unexpected SD genes, such as the interferon-related *sdY* in salmonids (Yano et al., 2013). The recent identification of several master SD genes in this group has been associated with the highly contiguous and reliable chromosome-level genome assemblies achieved via improvements in long-read sequencing technologies, scaffolding and bioinformatic approaches (Ramos and Antunes, 2022).

Flatfish (Pleuronectiformes) represent an emblematic fish order with notable adaptations to demersal life (Chen et al., 2014; Figueras et al., 2016; Robledo et al., 2017; Shao et al., 2017; Lü et al., 2021). They experience a remarkable metamorphosis from the bilateral morphology of pelagic larvae to the flat morphology typical of this group (Shao et al., 2017; Lü et al., 2021). The adaptation to demersal lifestyle promoted a quick evolutionary radiation, reflected by a higher molecular evolutionary rate than their bilateral counterparts (Lü et al., 2021). This rapid diversification has also led to a large variety of SD mechanisms, with diverse master SD genes and non-orthologous SD regions in all species analysed to date (Luckenbach et al., 2011; Martínez et al., 2021). The genomes of flatfish are compact (∼500-700 Mb; Robledo et al., 2017; Lü et al., 2021), which has facilitated the achievement of chromosome-level genome assemblies in species from several families of the order (Guerrero-Cózar et al., 2021; Lü et al., 2021).

The Senegalese sole (*Solea senegalensis*) is a valuable commercial flatfish with an expanding aquaculture industry (currently ∼2000 tons; Ana Riaza, pers. comm.). A number of genomic resources and tools have been generated for this species in the last decade, such as a comprehensive transcriptome and a derived oligo-microarray, several genetic maps, and recently, a whole genome assembly (Manchado et al., 2016; Guerrero-Cózar et al., 2021). As in several other flatfish species, Senegalese sole females outgrow males (Viñas et al., 2013), so obtaining all-female populations is an appealing strategy to increase growth rate. An important limitation for the expansion of Senegalese sole aquaculture is related to the low performance of F1 males (born and reared in captivity) compared to wild specimens reared in captivity, hampering the development of breeding programs in this species (Martín et al., 2019). Understanding the mechanisms underlying reproduction, including the SD mechanism, and how external cues are connected to gonad development via neural communication, will be essential to enable the growth of Senegalese sole aquaculture.

Here, we constructed a new highly contiguous genome assembly in Senegalese sole that we used to delve into the orthologous relationships of Pleuronectiformes by analysing the six most relevant commercial flatfish with chromosome-level genome assemblies pertaining to five different families. The reference assembly was used to re-sequence a sample of males and females which allowed the identification of the putative master SD gene of the species, the follicle stimulating hormone receptor (*fshr*), and confirmed an XX / XY system. Functional differences between males and females were evaluated throughout gonad development, from undifferentiated fry to adulthood. Our data suggest that an active Y-linked allele of the follicle stimulating hormone receptor (*fshry*) determines the testis fate for the undifferentiated primordium.

## Materials and Methods

### Genome assembly

Before assembly, reads were preprocessed as follows: the Illumina reads were trimmed using Trim-galore v0.6.6 (with options *--gzip -q 20 --paired --retain_unpaired*) (https://www.bioinformatics.babraham.ac.uk/projects/trim_galore/) and the nanopore reads were filtered using FiltLong v0.2.0 (with options *--min_length* 5000 *-- target_bases* 40,000,000,000) (FiltLong: https://github.com/rrwick/Filtlong). The filtering of nanopore data ensured having reads of at least 5 kb while optimizing for both length and higher mean base qualities, keeping 40 Gb (∼ 65x coverage).

We assembled the filtered ONT reads with NextDenovo v2.4.0 (https://github.com/Nextomics/NextDenovo) applying the options: minimap2_options_raw = -x ava-ont, minimap2_options_cns = -x ava-ont -k17 –w17 and seed_cutoff=10k. The resulting contigs were polished with Nextpolish v1.3.1 (Hu et al., 2020) using two rounds of long-read polishing and two rounds of short-read polishing. This assembly (fSolSen1.1) was evaluated with BUSCO v4.0.6 (Simão et al., 2015) using vertebrata_odb10, Merqury v 1.1 (Rhie et al., 2020) and Nseries.pl (an in-house script) to compute its contiguity. The Snailplot was created using blobtoolkit (Challis et al., 2020).

### Genome annotation

Repeats present in the fSolSen1 genome assembly were annotated with RepeatMasker v4-0-7 (http://www.repeatmasker.org) using the custom repeat library available for *Danio rerio*. Moreover, a new repeat library specific for our assembly was made with RepeatModeler v1.0.11. After excluding those repeats that were part of repetitive protein families (performing a BLAST search against UniProt) from the resulting library, RepeatMasker was run again with this new library in order to annotate the specific repeats.

The gene annotation of the assembly was obtained by combining transcript alignments, protein alignments and *ab initio* gene predictions. Firstly, RNAseq reads obtained from several tissues and developmental stages, either sequenced specifically in this study (brain, liver, bone, muscle, head kidney, fin, and 30 days post fertilization larvae) or existing in public databases, were aligned to the genome with STAR (Dobin et al., 2013) (v-2.7.2a). Transcript models were subsequently generated using Stringtie (Pertea et al., 2015) (v2.0.1) on each BAM file and then all the models produced were combined using TACO v0.6.2. Finally, PASA assemblies were produced with PASA (Haas et al., 2008) (v2.4.1). The *TransDecoder* program, which is part of the PASA package, was run on the PASA assemblies to detect coding regions in the transcripts. Secondly, the complete *Danio rerio, Scophthalmus maximus* and *Cynoglossus semilaevis* proteomes were downloaded from UniProt in April 2020 and aligned to the genome using SPALN (Slater & Birney, 2005) (v2.4.03). *Ab initio* gene predictions were performed on the repeat-masked fSolSen1 assembly with three different programs: GeneID (Parra et al., 2000) v1.4, Augustus (Stanke et al., 2006)□v3.3.4 and Genemark-ES (Lomsadze et al., 2014) v2.3e with and without incorporating evidence from the RNAseq data. The gene predictors were run with trained parameters for human, except Genemark that runs in a self-trained manner. Finally, all the data was combined into□consensus CDS models using EvidenceModeler-1.1.1 (EVM, Haas et al., 2008). Additionally, UTRs were identified and alternative splicing forms annotated through two rounds of PASA annotation updates. Functional annotation was performed on the annotated proteins with Blast2go (Conesa et al., 2005). First, a Diamond blastp (Buchfink et al., 2021) search was made against the nr (last accessed May 2021) and Uniprot (last accessed August 2021) databases. Furthermore, Interproscan (Jones et al., 2014) was run to detect protein domains on the annotated proteins. All these data were combined by Blast2go, which produced the final functional annotation results.

The non-coding RNA annotation required several steps. First, we annotated as long-non-coding RNAs (lncRNAs) those PASA assemblies that had not been included into the protein-coding annotation, that did not match any protein-coding gene and that were longer than 200bp.

We also sequenced small RNAs (sRNAs) from several tissues and developmental stages, including bone, muscle and fin. The corresponding reads were aligned with STAR [1] (v-2.7.2a) with parameters (-outFilterMultimapNmax 25 --alignIntronMax 1 - -alignMatesGapMax 1000000 --outFilterMismatchNoverLmax 0.05 -- outFilterMatchNmin 16 --outFilterScoreMinOverLread 0 -- outFilterMatchNminOverLread 0). The resulting mappings were processed to produce the annotation of small non-coding RNAs. First, TACO was run to assemble the reads into transcripts. Transcripts overlapping exons from the protein-coding or lncRNA annotations were removed from the set of small non-coding RNAs. Finally, the program cmsearch (Cui et al., 2016) (v1.1.4) that comes with Infernal (Nawrocki & Eddy, 2013) was run on the sncRNAs against the RFAM (Nawrocki et al., 2015) database of RNA families (v14.6) in order to annotate products of those genes.

The final non-coding annotation contains the lncRNAs and the sncRNAs. The resulting transcripts were clustered into genes using shared splice sites or significant sequence overlap as criteria for designation as the same gene.

The final non-coding annotation contains the lncRNAs and the sncRNAs. The resulting transcripts were clustered into genes using shared splice sites or substantial sequence overlap as criteria for designation as the same gene.

### Genetic map construction and genome scaffolding

Six Senegalese sole broodstock, three males and three females, were used for founding three full-sib families using *in vitro* fertilisations as described for the Proof-of-concept experiment by Ramos-Júdez et al. (2021). Briefly, females were selected by maturity status and induced to ovulate with an injection of 5 μg kg^-1^ of GnRHa (Sigma code L4513, Sigma, Spain), whilst no hormones were used on males. Gametes were extracted from females and males using gentle abdominal pressure, fertilised (1 male x 1 female) and incubated in 30 L incubators. Larvae were randomly sampled from the incubators and placed in 70% ethanol six days after hatching. All this work was performed at the Sant Carles de la Rápita Center (Catalonia, Spain).

Library preparation followed the 2b-RAD protocol (Wang et al., 2012) with slight modifications (Maroso et al., 2018). Briefly, DNA samples were adjusted to 80 ng/μL and digested using the IIb type restriction enzyme AlfI (Thermo Fisher). Specific adaptors and individual sample barcodes were ligated and the resulting fragments amplified. After PCR purification, samples were quantified and equimolarly pooled. The pools were sequenced on a NextSeq 500 Illumina sequencer in the FISABIO facilities (Valencia, Spain). A total of 81, 77 and 71 offspring from the three full-sib families founded (F1, F2 and F3, respectively) were evenly mixed and sequenced in three independent runs with parents also included at double concentration.

Demultiplexed reads according to the sample barcodes were first trimmed to 36 nucleotides and a custom perl script was used to remove reads without the AlfI recognition site in the correct position. Then, reads were processed using the *process_radtags* module in STACKS v2.0 (Catchen et al. 2013), removing reads with uncalled nucleotides or a mean quality score below 20 in a sliding window of 9 nucleotides. Bowtie 1.1.2 (Langmead et al. 2009) was used to align the filtered reads against the assembled genome (see above), allowing a maximum of three mismatches and a unique valid alignment. Finally, the output files were used to feed the *gstacks* module in STACKS, using the *marukilow* model to call variants and genotypes.

To build the genetic map, SNPs with extreme deviations from Mendelian segregation (P < 0.001) were removed, and only informative SNPs genotyped in at least 60% offspring were used. The *grouping* function of JoinMap 4.1 (Stam, 1993) was used to build linkage groups (LG), based on an increasing series of LOD scores from 7.0 to 10.0 to accommodate to the 21 chromosomes (C) of the Senegalese sole karyotype (Vega et al., 2002). Marker ordering was performed using the Maximum Likelihood (ML) algorithm with default parameters with the Kosambi mapping function used to compute centi-Morgans (cM) map distances. Map consensus were built using MergeMap (Wu et al., 2011) and visualized with MapChart 2.3 (Voorrips 2002) and Circos software (Krzywinski et al., 2009).

Chromonomer 1.13 (Catchen et al., 2020) was used to anchor and orient the contigs of the genome to the genetic map. Further, the correspondence between genetic map and contigs enabled the manual curation of original contigs that mapped to different linkage groups (LG). This occurred with the longest contig (see Results), which was split into into two fragments at a specific position, determined by comparing the sequence of the contig located between the closest markers mapping in different linkage groups with orthologous regions of other flatfish chromosome-level genomes available in Ensembl. The program RIdeogram was used to visualize the correspondence between genome contigs and crhomosomes (Hao et al., 2020).

### Cytogenetic map and mapping integration

The BAC clones used for mapping came from a *S. senegalensis* BAC library including 29,184 clones. We used the methodology for BAC identification through 4D-PCR and DNA purification described by García-Cegarra et al. (2013). BAC clones were sequenced using either a Genome Sequencing FLX System or MiSeq Illumina by the LifesequencingTM Platform (Valencia, Spain). Assembly, functional and structural annotation of BAC sequences was carried out as previously reported (Merlo et al., 2021).

Chromosome preparations were obtained from colchicine-treated larvae according to García-Cegarra et al. (2013) and from spleen and anterior kidney culture according to the following procedure. Adult fish were anesthetized with clove oil (40 mg/L), then injected intraperitoneally with colchicine (0.05 %) and kept in an oxygenated tank for 3–4 h. A clove oil overdose was used to kill the individuals and spleen and anterior kidney were extracted and dispersed in a 0.056 % KCl solution. The solution was filtered (mesh size 40 to 100 μm), subjected to a hypotonic shock with KCl and fixed in Carnoy solution. Positive BAC clones were isolated following Merlo et al. (2021) and labelled either by nick translation (García-Cegarra et al., 2013) or by DOP-PCR (Portela-Bens et al., 2017). Pre-treatment, hybridization, post-hybridization and staining for double-fluorescence in situ hybridization (FISH) followed García-Cegarra et al. (2013) and for multiple-FISH Portela-Bens et al. (2017). To capture double- and multiple FISH images the fluorescence microscope Zeiss PALM MicroBeam equipped with an AxioCam MRm digital camera and a digital CCD camera (Olympus DP70) coupled to a fluorescence microscope (Olympus BX51 and/or Zeiss Axioplan using software of MetaSystems, Altlussheim, Germany), respectively, were used. Experimental procedures were carried out following the regulations of the University of Cádiz (Spain) for the use of laboratory animals and the Guidelines of the European Union Council (86/609/EU).

To establish the correspondence between LGs / genome scaffolds and the chromosomes for mapping integration, ∼141 BAC clones were positioned on chromosomes using BAC-FISH. Then, BAC sequences were located in the 21 scaffolds of the genome by a megablast search tool from blast algorithm (Altschul et al., 1990) using the following parameters: Evalue < E-20; max_hsps = 10; sequence overlap > 5 kb. Alignments were visualized with the Integrative Genomics Viewer (IGV) program (Robinson et a., 2011) and manually explored.

### Comparative mapping: orthology with other flatfish

We took advantage of the recent enrichment of Pleuronectiformes genomic resources, to investigate orthology within this emblematic and controversial fish group (Figueras et al., 2016; Robledo et al., 2017). We chose six commercially relevant flatfish species with genomes assembled at chromosome-level and annotated proteomes (NCBI database, July 2021), representing five different families, and including the *S. senegalensis* genome here assembled and annotated, pertaining to the family Soleidae. The other five proteomes corresponded to *C. semilaevis* (Cynoglossidae), *Hippoglossus hippoglossus* and *H. stenolepis* (Pleuronectidae), *Paralichthys olivaceus (*Paralychthydae) and *S. maximus* (Scophthalmidae) (Bioprojects PRJNA251742, PRJNA562001, PRJNA622249, PRJNA73673 and PRJNA631898, respectively). For each proteome, the longest isoform of each protein was filtered using CD-HIT (Li & Godzik, 2006) and used for the analysis with OrthoFinder (Emms & Kelly, 2019), to infer orthogroups, orthologs, gene trees and the rooted species tree, using the reference proteome of *D. rerio* (Bioproject PRJNA11776). The shared set of single-copy orthologs anchored to chromosomes in the six species studied was used for species phylogenetic tree construction and for comparative mapping. The *P. olivaceous* gene sequences were mapped onto the chromosome-level genome assembly v1.0 of the species (GCA_001970005.2; Bioproject PRJNA344006) using BLAST (Altschul et al. 1990). The syntenic relationships among the chromosomes of the six flatfish species were represented using ggsankey (https://github.com/davidsjoberg/ggsankey), while syntenic comparisons by pairs were drawn using the R package circlize (Gu et al, 2014). Finally, we compared the Senegalese sole genome here reported with the previous one by Guerrero-Cozar et al. (2021) using LASTZ (Harris, 2007) (options “--notransition -- step=400 --nogapped --format=rdotplot”).

### Sex determining (SD) gene candidates

Six adult males and six adult females were re-sequenced using 150 bp PE reads on an Illumina NovaSeq 6000 System to 20x coverage in the CNAG Platform and aligned independently against our newly assembled reference Senegalese sole genome. A broad SNP dataset was identified and genotyped in those twelve males and females using SAMtools and filtered according to default parameters. This SNP dataset was used to estimate the relative component of genetic differentiation between males and females (F_ST_) and the intra-sex fixation index (F_IS_) across the whole genome using GENEPOP 4.7.5 (Raymond & Rousset, 1995). F_ST_ and F_IS_ values were averaged over 50 consecutive SNPs and explored using sliding windows across each chromosome of the genome to look for deviation from the null hypothesis (F_ST_ and F_IS_ = 0).

### Developing a tool for sexing

As shown in Results, *fshr* was the most consistent sex determining (SD) candidate gene. It exhibited a large number of diagnostic markers across its whole length, homozygous in females and heterozygous in males, which agrees with the XX / XY system reported for Senegalese sole (Molina-Luzón et al., 2015). Several of these markers were used to develop a molecular tool to identify sex using a non-invasive method (e.g. from a fin-clip). This tool was very valuable to evaluate gene expression across gonad development from the undifferentiated germinal primordium, specially at those stages where the gonad was still undifferentiated (see below). A SNaPshot assay was developed for three diagnostic markers, chosen by their technical feasibility, i.e. with no other polymorphism within ±100 bp from the SNP, according to the resequencing information of six males and six females. Two external primers to amplify a surrounding region of ∼150 bp and one internal adjacent to the SNP for the mini-sequencing reaction were designed using Primer 3 (Rozen & Skaletsky 2000). PCR was performed on a Verity ™ 96-Well Thermal Cycler (Applied Biosystems) as follows: initial denaturation at 95 °C for 5 min, 30 cycles of denaturation at 94 °C for 45 s, annealing temperature at 58°C for 50 s, and extension at 72 °C for 50 s; a final extension step was done at 72 °C for 10 min. Subsequently, 1 μL of the PCR product was purified by incubation with 0.5 μL illustra™ ExoProStar™ 1-STEP Kit at 37 °C for 15 min followed by 85 °C for 15 min to eliminate unincorporated primers and dNTPs. The SNaPshot minisequencing reaction was carried out using the SNaPshot® Multiplex Kit (Applied Biosystems) in an ABI Prism 3730xl DNA sequencer. For each reaction, 1.5 μL of purified PCR product, 0.5 μL (2 μM) of internal primer and 2 μL of SNaPshot™ Multiplex Ready Reaction Mix were used in a final volume of 5 μL. The reaction profile consisted initial denaturation at 96 °C for 1 min, 30 cycles at 96 °C for 10 s, 55 °C for 5 s, and 60 °C for 30 s. The extension product was incubated with 1 μL of shrimp alkaline phosphatase at 37 °C for 60 min followed by 85 °C for 15 min to remove unincorporated dideoxinucleotides (ddNTPs) after thermal cycling.

### Gonad differentiation: histological evaluation

Five fishes per sex were collected from each stage at 84, 98 and 126 days post fertilization (dpf), juveniles (315 dpf) and mature adults (810 dpf) at Stolt Sea Farm SL facilities (Ribeira / Cervo, Spain) and sacrificed by decapitation. All fishes were maintained at the same standard temperature, air-flow and feeding conditions of the usual production protocol of the company until sacrifice. Gonads were dissected fresh for macroscopic evaluation and classified as testes or ovaries by visual inspection in juveniles and adults. At the three initial stages, the molecular tool outlined before was used for sexing. Then, the gonads were fixed by immersion in 4% paraformaldehyde and later embedded in paraffin wax to be cut into sagital sections 3-6 μm and stained with haematoxylin-eosin for optical microscopy evaluation. Experimental procedures for the use of farm animals were carried out following the regulations of the University of Santiago de Compostela and Stolt Sea Farm SA company (Spain) and the Guidelines of the European Union Council (86/609/EU).

### Gonad differentiation: gene expression

#### qPCR

Gene expression of the candidate gene along with several genes markers of the initial stages of gonadal differentiation were evaluated through qPCR on five male and five female gonads at the five developmental stages considered: 84, 98 and 126 dpf, juveniles and adults. RNA extraction was performed using the RNeasy mini kit (Qiagen) with DNase treatment and RNA quality and quantity were evaluated in a Bioanalyzer (Bonsai Technologies) and in a NanoDrop® ND-1000 spectrophotometer (NanoDrop® Technologies Inc), respectively. Primers for qPCR of the candidate gene (*fshr*, see Results), and for ovary (*cyp19a1*, aromatase) and testes (*amh*, anti-mullerian hormone; *sox9*, SRY-Box Transcription Factor 9) markers (Robledo et al., 2015), along with those related to germinal cell proliferation (*gsdf*, gonadal soma-derived factor, and *vasa*, ATP-dependent RNA helicase) were designed using the Primer 3 software. Reactions were performed using a qPCR Master Mix Plus for SYBR Green I No ROX (Eurogenetec) following the manufacturer instructions, and qPCR was carried out on a MX3005P (Agilent Technologies). Analyses were performed using the MxPro software (Agilent). The ΔΔCT method was used to estimate expression taking the ribosomal proteins S4 (*rps4*) and L17 (*rpl17*) genes, and ubiquitin (*ubq*) as reference genes. These three genes had been previously validated for qPCR in turbot gonads by Robledo et al. (2014). Two technical replicates were included for each sample. T-tests were used to determine significant differences between sex. Additionally, preliminary data from an RNAseq study on gonad differentiation was specifically addressed on the SD candidate *fshr* gene (see Results) using diagnostic SNPs on exons, to ascertain which allele of the gene, X- or Y-linked, was expressed at the different stages of gonadal differentiation in males and females, and to identify splicing variants that could be involved in triggering the SD signal.

## Results

### Genome assembly and annotation

The initial assembly comprised 82 contigs with an N50 of 23.4 Mb (sizes ranging from 0.3 to 30.1 Mb) for a total assembly size of 613 Mb (Table S1). A high-density genetic map was used to place 51 contigs, representing 99.0% of the whole assembly (607.9 Mb), into the 21 chromosomes of the Senegalese sole haploid karyotype (n = 21; Vega et al., 2002) (see below). This assembly is highly contiguous (23.4 Mb contig N50, 29.0 Mb scaffold N50), and showed high consensus quality (QV = 43.17) and gene (97.8% Complete BUSCO genes) and k-mer (98.18%) completeness (Table S1). Consistent with the low percentage of duplicated BUSCOs (0.6%), the k-mer spectra (Fig. S1) did not reveal any evidence of artificial duplications. The repetitive peak observed at 360x very likely corresponds to true long repetitive regions captured by the nanopore reads (Fig. S1).

Repetitive sequences made up to 8% of the Senegalese sole genome (Table S2). These were grouped into three main categories: simple repeats (2.81%), low-complexity motifs (0.32%), and transposable elements (TEs) (4.73%). The TE-derived fraction was very similar to that found in other high-quality flatfish genome assemblies (5.8% and 5% in *C. semilaevis* and *S. maximus*, respectively). The Senegalese sole genome displayed a higher TE proportion than *T. nigroviridis* and *Fugu rubripes* (< 3%), but much lower than that observed in other fish such as *Danio rerio* (> 40%) (Gao et al., 2016).

In total, 24,264 protein-coding genes producing 40,511 transcripts (1,67 transcripts per gene) and encoding for 37,259 unique protein products, were annotated in the Senegalese sole genome (Table S3). We were able to assign functional labels to 85% of the annotated proteins. The annotated transcripts contained 12.79 exons on average, with 95% of them being multi-exonic. In addition, 52,888 non-coding RNAs were annotated (6,871 long non-coding and 46,017 small RNAs).

### Genetic map construction and genome scaffolding

RAD-seq of three full-sib Senegalese sole families, consisting of 81 (F1), 77 (F2) and 71 offspring (F3), and the six correspondent parents, yielded around one billion raw reads, representing on average 11.9 million reads per parent and 4.2 million per offspring (Table S4). After filtering, 82.9% reads were retained and aligned to the Senegalese sole genome. Most of the filtered reads were correctly aligned to the reference genome according to the parameters established (91.3%), resulting in ∼9.5 million reads per parent and ∼3.1 million per offspring.

The *gstacks* module rendered a total of 156,981 loci, 35,441 of them containing at least one single nucleotide polymorphism (SNP). Applying the mapping filtering criteria, 16,890, 15,457 and 16,715 SNPs from F1, F2 and F3, respectively, were retained for map construction. From these, 4889 SNPs were shared among the three families, resulting in a total of 29,126 unique SNPs for mapping, which represent 47.5 markers per Mb or 1 SNP every 21.1 kb.

Separate male and female genetic maps were built in each family. A strict LOD score of 9.0 was applied to achieve a consistent number of LGs across all maps matching the n = 21 chromosomes of the species (Vega et al., 2002). The average number of markers per LG across all maps was 434 (ranging from 93 to 694 markers) (Table S5 and S6; Figure 1). Female maps were slightly longer than male maps, with an average F *vs* M recombination ratio of 1.1:1. The 4889 shared informative markers across families were used to establish their correspondence and build the female, male, and species consensus maps. The final species consensus map included 28,838 markers across 21 LGs embracing 40,704.47 cM (Table S5 and S6; Figure 1).

**Figure 1:**
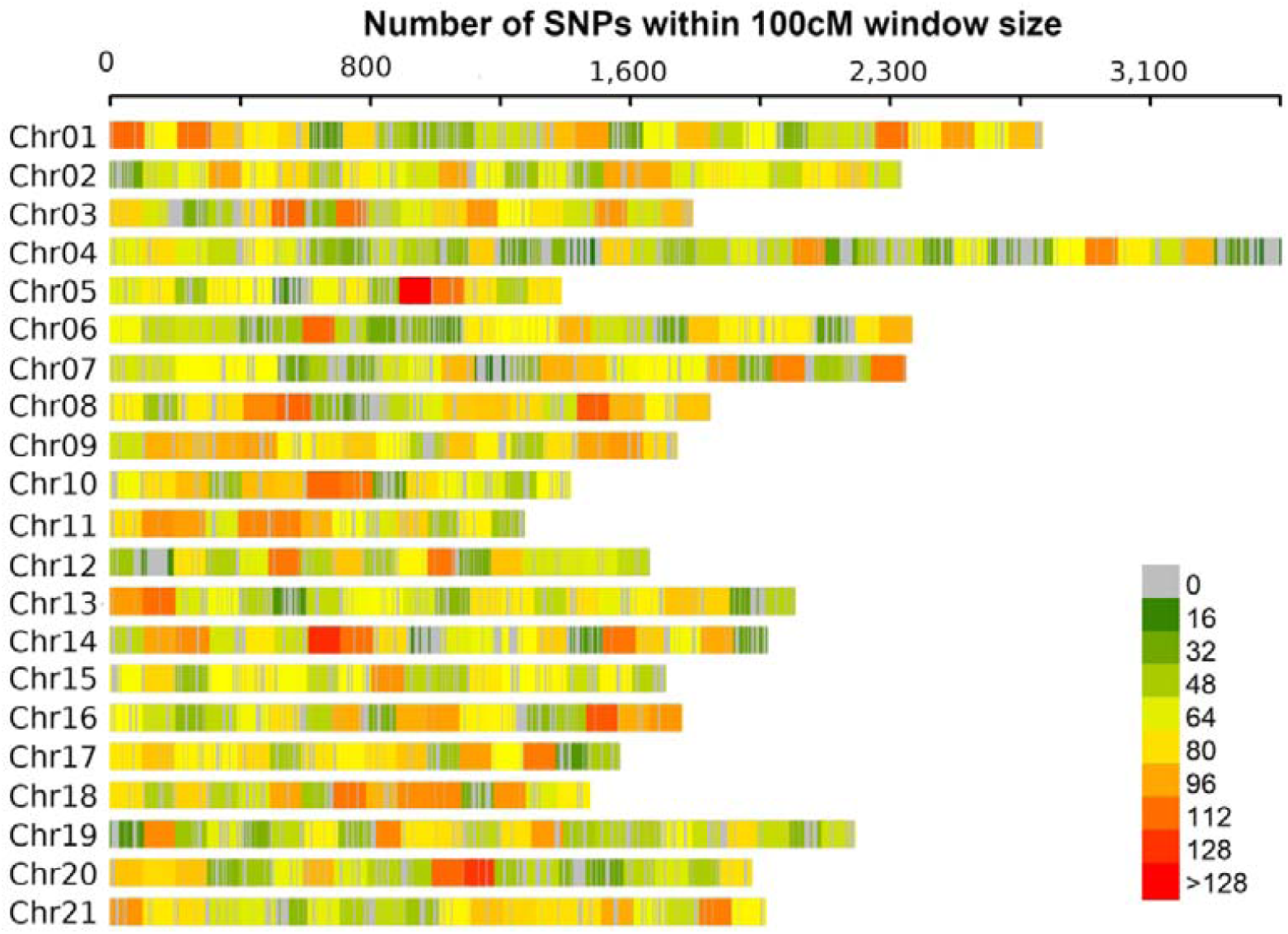
Mapping marker density across linkage groups / chromosomes in the *Solea senegalensis* consensus map

The consensus genetic map was used to anchor 51 contigs into the 21 *S. senegalensis* scaffolds as described above. The Chromonomer program anchored and oriented a total of 51 scaffolds to the consensus map (Table S7) and only one contig was split into two fragments assigned to LG6 and LG9 (Figure 2). This resulted in 99.02% of the 613 Mb placed into the 21 chromosomes of the Senegalese sole karyotype, improving upon the 90.0% reported by Guerrero-Cózar et al. (2021) and similar to other flatfish species recently assembled following a similar methodology (Einfeldt et al., 2021; Zhao et al., 2021; Ferchaud et al., 2022) (Table S8). The new assembly showed a high one-to-one correspondence at the chromosome level with the previous version of Guerrero-Cózar et al. (2021) (Fig. S2; Table S5). However, the substantial fragmentation of the previous version (1937 scaffolds *vs* 82 contigs) gave rise to discrepancies related to wrong orientation of many minor contigs within many of the chromosomes, especially the smaller ones (e. g. C14, C15, C17, C18).

**Figure 2:**
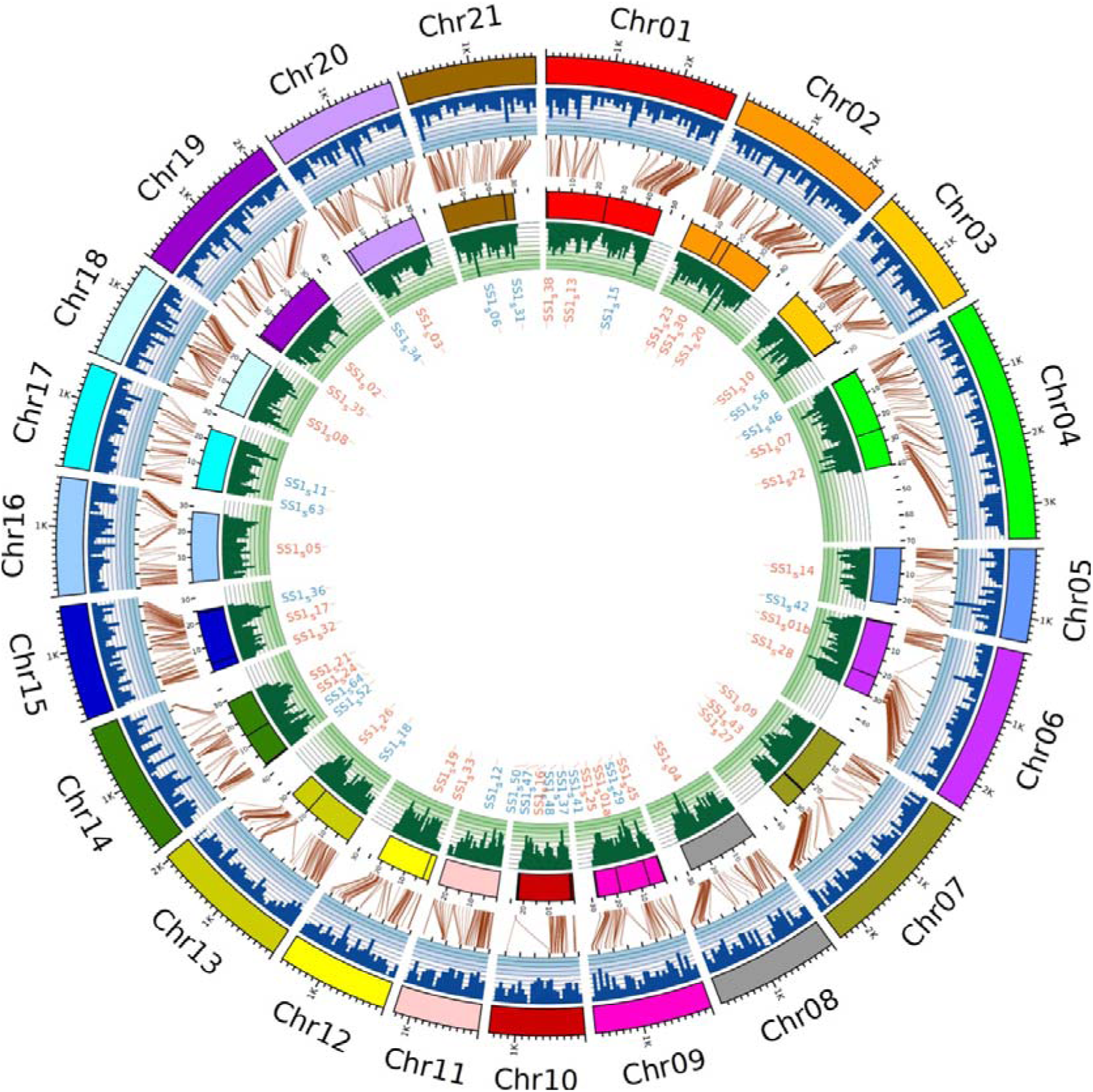
Circos plot of the genome map and anchored scaffolds in *Solea senegalensis*. From outer to inner circles are represented: the 21 LGs / Chromosomes (tick marks every 100 cM); histograms of the number of markers per 50 cM (in dark blue); brown lines indicating the position of the collinear markers selected by Chromonomer in the genetic map and in the genome scaffolds; the 51 anchored scaffolds (tick marks every 5 Mb); histograms of the number of markers per Mb (in dark green); and the names of the scaffolds (in red those anchored in reverse strand).

### Cytogenetic map and mapping integration

A total of 141 BACs were used to anchor the LGs / scaffolds to the chromosomes of *S. senegalensis* karyotype using BAC-FISH information from previous reports and from this study (Table S9; Figs. S3A and S3B). On average 6.6 BACs per scaffold were used to establish the correspondence between the genetic, physical and cytogenetic maps (range: 4 to 14 BACs), thus providing a better, more robust mapping integration within this study (Figure 1) with respect to the previous assembly by Guerrero-Cózar et al. (2021). Ten out of 141 BACs hybridized to more than one chromosome or to different locations within the same chromosome suggesting paralogous regions in the genome (Table S9). The minor 5S rDNA was located on C6 and C11, while signals of the major rDNA (18S + ITS1 + 5,8) were found at C6 and C20.

### Comparative mapping: orthology with other flatfish

The new Senegalese sole assembly was compared with the chromosome-level genomes of five other commercial flatfish, representing in total five different Pleuronectiformes families. From 157,596 gene sequences analysed in the six species, 151,659 (96.2%) were assigned to 21,367 orthogroups, 16,522 with genes in the six species, and 7936 corresponding to single copy orthogroups (Tables S10 and S11A). Gene trees were obtained for all orthogroups with more than four sequences from two or more species, along with the unrooted species tree (Figs. S4A, S4B, S4C). Nearly all single-copy orthologs detected in the six flatfish species (99.9%) mapped to the Senegalese sole genome, and in the other species this figure ranged from 97.4% in the tongue sole to 100% in the turbot (Table S11A), supporting the completeness of these flatfish genomes. A rooted tree based on 5,870 conserved single-copy orthologs between flatfish and zebrafish reconstructed the known flatfish phylogeny (Table S10; Table S11B; Fig. S4D).

A total of 7,603 single-copy orthologs (95.8%) were anchored to the chromosomes of the six species and used for comparative mapping (Figure 3; Table S11A). Most *S. senegalensis* chromosomes showed highly conserved macrosyntenic patterns with other flatfish chromosomes (Figure 3; Fig. S5; Table S11 and S12). Major one-to-one chromosome orthology was observed between *P. olivaceus* (Pol), *H. stenolepis* (Hst) and *H. hippoglosus* (Hhi), pertaining to Pleuronectidae and Paralychthydae, families that exhibit the ancestral chromosome number of the order (n = 24; Pardo et al., 2001). More complex chromosomal reorganizations were observed in the more divergent karyotypes of *S. senegalensis* (Sse; n=21), *C. semilaevis* (Cse; n=21) and *S. maximus* (Sma; n=22) (Figure 3; Tables S11A and S12), involving several biarmed chromosomes, such as those observed in the Senegalese sole karyotype (Vega et al., 2002).

**Figure 3:**
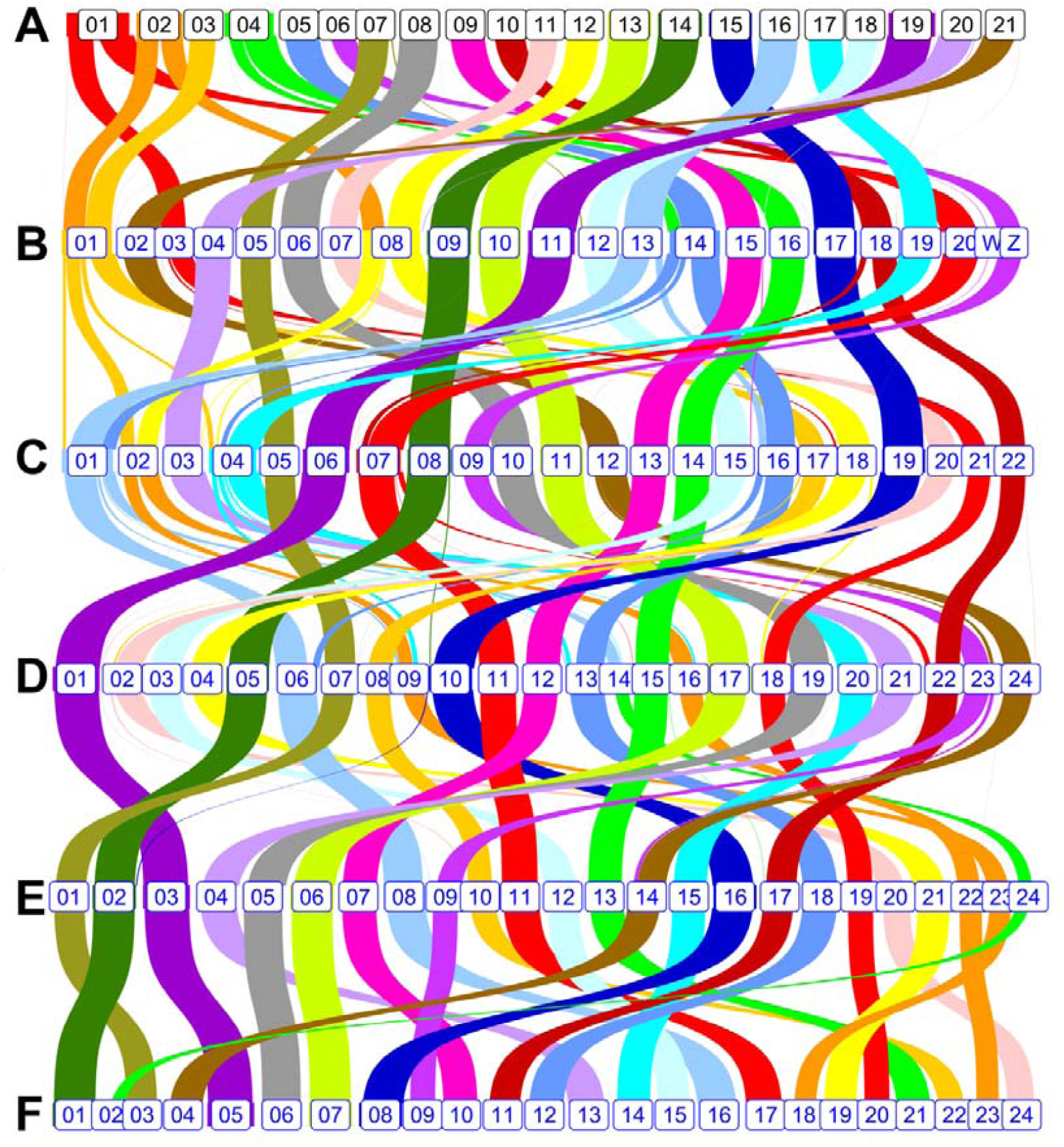
Macrosyntenic conservation of six chromosome-level flatfish genomes. The number in the squares represents the chromosome identity for each species: A) *Solea senegalensis* (Family Soleidae), B) *Cynoglossus semilaevis* (Cynoglossidae), C) *Scophthalmus maximus* (Scophthalmidae), D) *Paralichthys olivaceus* (Paralychthydae) E) *Hippoglossus stenolepis* (Pleuronectidae), F) *Hippoglossus hippoglossus* (Pleuronectidae).

Consistent macrosyntenies involving orthology of two acrocentric chromosomes across all species were observed with respect to both arms for the largest metacentric Sse C1 (Cse20-Sma7-Pol11-Hst11-Hhi17 and Cse3-Sma21-Pol18-Hst19-Hhi20) and for the submetacentric Sse C4 (Cse14-Sma1-Pol14-Hst24-Hhi2 and Cse16-Sma14-Pol15-Hst13-Hhi21), suggesting centric fusions in the Senegalese sole lineage. The second largest metacentric Sse C2 also shared major syntenic blocks with two acrocentric chromosomes in most flatfish (Cse1-Pol9-Hst23-Hhi18; Cse8-Pol16-Hst22-Hhi23), but matched to a single metacentric in turbot (Sma2), suggesting independent fusions of the same chromosomes in two different lineages (Fig. S6). Other Senegalese sole biarmed chromosomes (Sse C3, C5-C9) matched each to single acrocentric chromosomes in all other flatfish, such as the metacentric Sse C3 (Cse1-Sma17-Pol8-Hst10-Hhi22; Fig. S7) or the submetacentric Sse C5 (Cse14-Sma16-Pol13-Hst18-Hhi12; not shown), consistent with pericentric inversions. The 12 acrocentric chromosomes of Senegalese sole (Sse C10-C21) all showed major one-to-one correspondence with uniarmed chromosomes across all flatfish lineages (Figure 3). Finally, fine comparative mapping showed extensive reorganizations at microsyntenic scale, suggesting that many inversions and intrachromosomic translocations should have taken place in the karyotype evolution of Pleuronectiformes in accordance with their phylogenetic distance (Table S12; Fig. S7 and S8).

### Sex determining (SD) gene candidate

Whole-genome resequencing of six males and six females identified a total of 9,078,413 SNPs. The relative component of genetic differentiation (F_ST_) between males and females and the intrapopulation fixation index in the whole sample (F_IS_) was estimated across the whole genome using a 50 SNP sliding window from genotyping data. A region at C12, between 11,700,000 and 11,736,000 bp and embracing 15,317 SNPs, showed a very consistent pattern of genetic differentiation between male and female samples (average F_ST_ = 0.304; range: from 0.105 to 0.476) and a significant heterozygote excess in the whole population (average F_IS_ = - 0.519; range from -0.041 to -0.903) (Table S13; Figure 4). This region included the 14 exons of the *fshr* gene (from 11,717,616 to 11,726,742 bp; Table S14). Out of 284 SNPs within the gene, 168 showed consistent differences between sexes (diagnostic markers), the males being always heterozygous and females homozygous according with the XX / XY system reported for this species (Molina-Luzón et al., 2015). A total of 33 diagnostic variants were located within exons of *fshr* (24 non-synonymous), 16 of them in exon 14 (11 non-synonymous). This asymmetry is consistent with the distribution of F_ST_ and F_IS_ values, more extreme towards the 3’ end of the gene (close to 0.5 and -1, respectively) (Figure 4). The accumulation of non-synonymous variants might suggest the degeneration of the Y-linked *fshr* gene, representing a non-functional variant.

**Figure 4:**
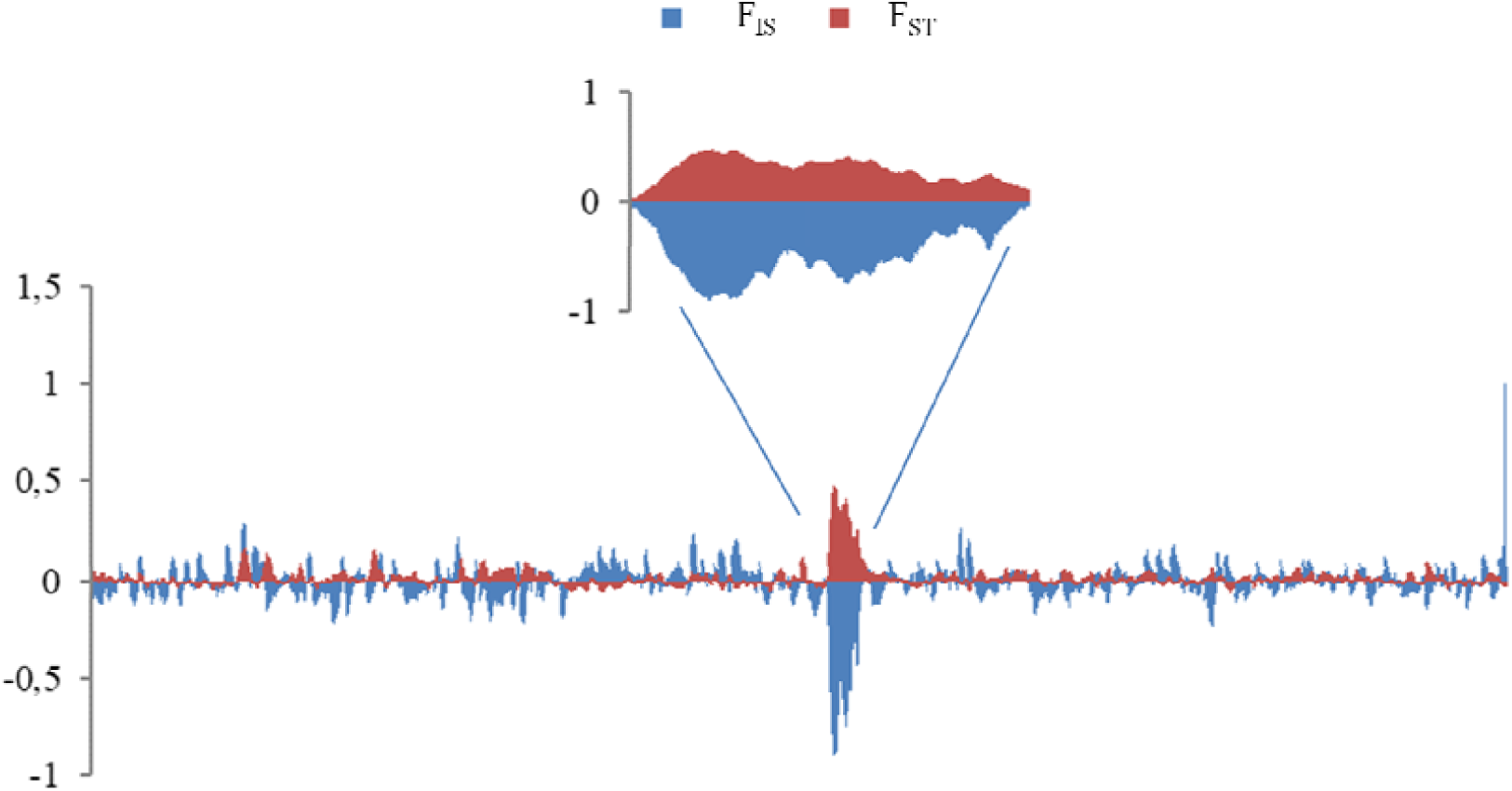
Genetic differentiation (F_ST_) and intrapopulation fixation index (F_IS_) between male and female populations at scaffold 19 (C12) of *Solea senegalensis*

### A molecular tool for sexing

Three diagnostic SNPs surrounded by ± 100 bp conserved regions (no polymorphisms) were selected for assessing genetic sex using a SnaPshot assay. Three sets of primers (two external and one internal) were designed (Table S15) and tested in five adult males and females. One marker differentiated males and females and matched the *in silico* genotype expectations, and was further validated in 48 males and 48 females pertaining to the broodstock of a Senegalese sole company. In all but three fishes, the genetic sex matched the phenotypic sex. Those three fishes were phenotypic males sexed as genetic females. Nonetheless, among a total of 133 fishes with known sex analysed in this study (see below RNAseq), only those three males showed a discordant genotype (2.3%).

### Gonad differentiation: gene expression analysis

#### qPCR

We analysed gene expression of *fshr* and other relevant genes (*amh, cyp19a1, gsdf, vasa*, and *sox9a*) related to gonadal differentiation, from the undifferentiated primordium until mature adults, including 86, 98 and 126 dpf, juveniles (308 dpf) and adults (810 dpf), using five males and five females at each stage, sexed either macroscopically or with the SS-sex marker at undifferentiated stages. Gonads of adults, juveniles and 128 dpf were macroscopically identified, while those from 98 dpf and, especially, 86 dpf were hard to identify (Fig. S9). Histological analyses of all stages were in accordance with previous information by Viñas et al. (2013) (Figs. S10 and S11), excluding 86 dpf, not previously reported, which showed an undifferentiated pattern similar to that observed at 98 dpf but smaller in size (Fig. S12).

Surprisingly, the *fshr* gene was significantly overexpressed in males at all stages (notice the putative aforementioned degeneration suggested for the Y-linked allele), extremely in juveniles, but even at the undifferentiated 84 dpf and 98 dpf stages (Figure 5), an observation corroborated by the RNAseq data (Fig. S13). RNAseq data were also used to check the expression of the X-linked and the Y-linked *fshr* alleles, taking advantage of the presence of diagnostic SNPs associated to the X and Y alleles. Interestingly, the Y-linked allele was expressed in males from 84 dpf to juveniles; furthermore, it showed higher expression than the X-linked allele across all stages (Wilcoxon rank pair-related sample test; P = 0). In adults of both sexes expression was nearly undetectable. Despite the low number of replicates and the substantial variation between them, differences were also significant at most stages: 98 dpf, 126 dpf and juveniles (P = 0.043), but even at 86 dpf, where *fshr* started to be active (three individuals out of five passing filtering genotyping parameters), the Y-linked allele showed 50% more counts on average than the X-linked (323.7 *vs* 203.7), although not significant (P = 0.102) (Figure 6; Table S16). The read counting across the 14 exons of *fshr* showed a very similar profile both in males and in females, and the correlation between X- and Y-linked alleles across exons in males was highly significant (*r* = 0.981; P = 0), suggesting no alternative splicing variants in the gene or in the two alleles at the evaluated stages (Fig. S14).

**Figure 5:**
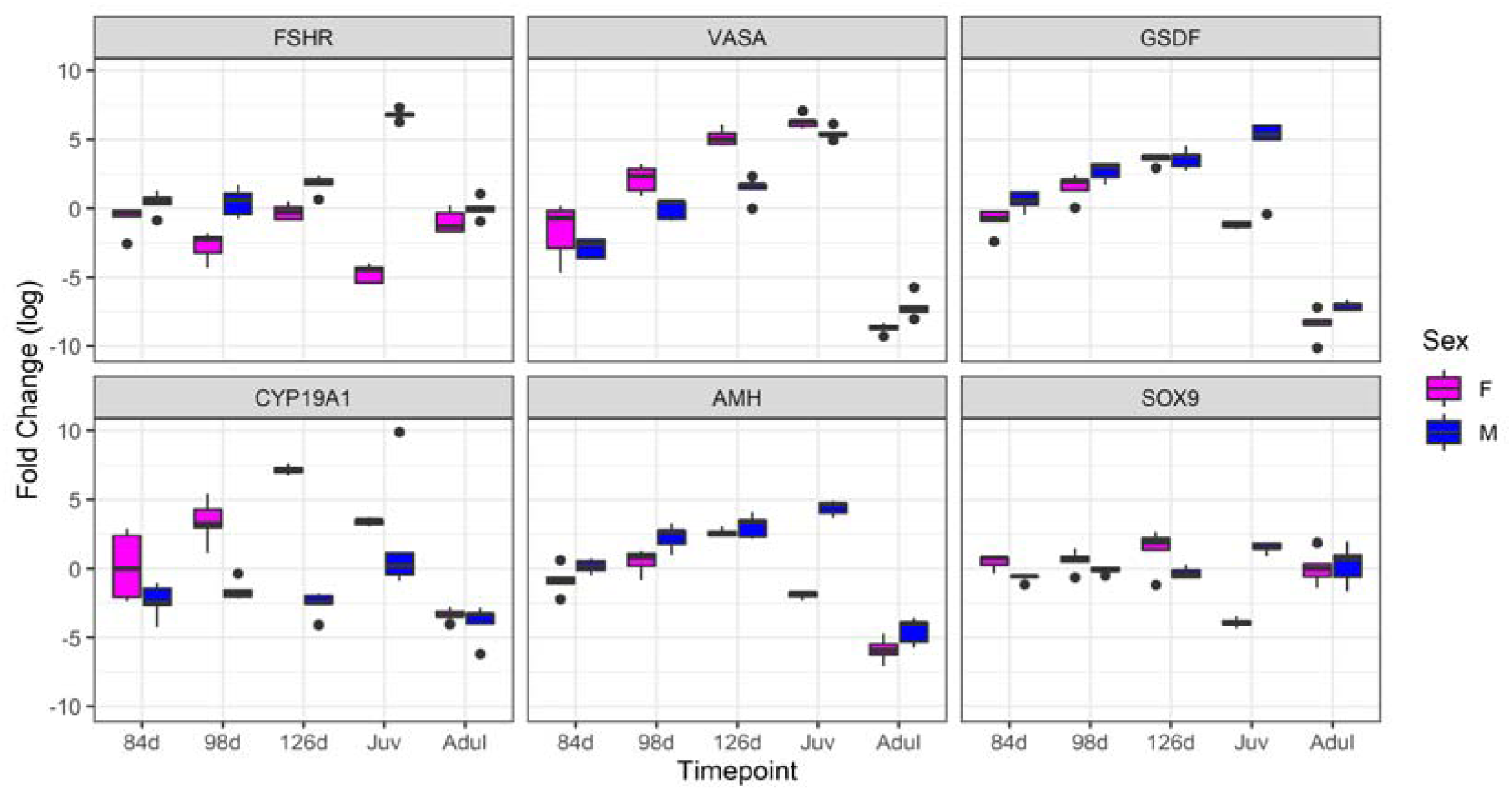
Box plots of the qPCR for the follicle stimulating hormone receptor SD gene of *Solea senegalensis* and other key marker genes across gonad development.

**Figure 6:**
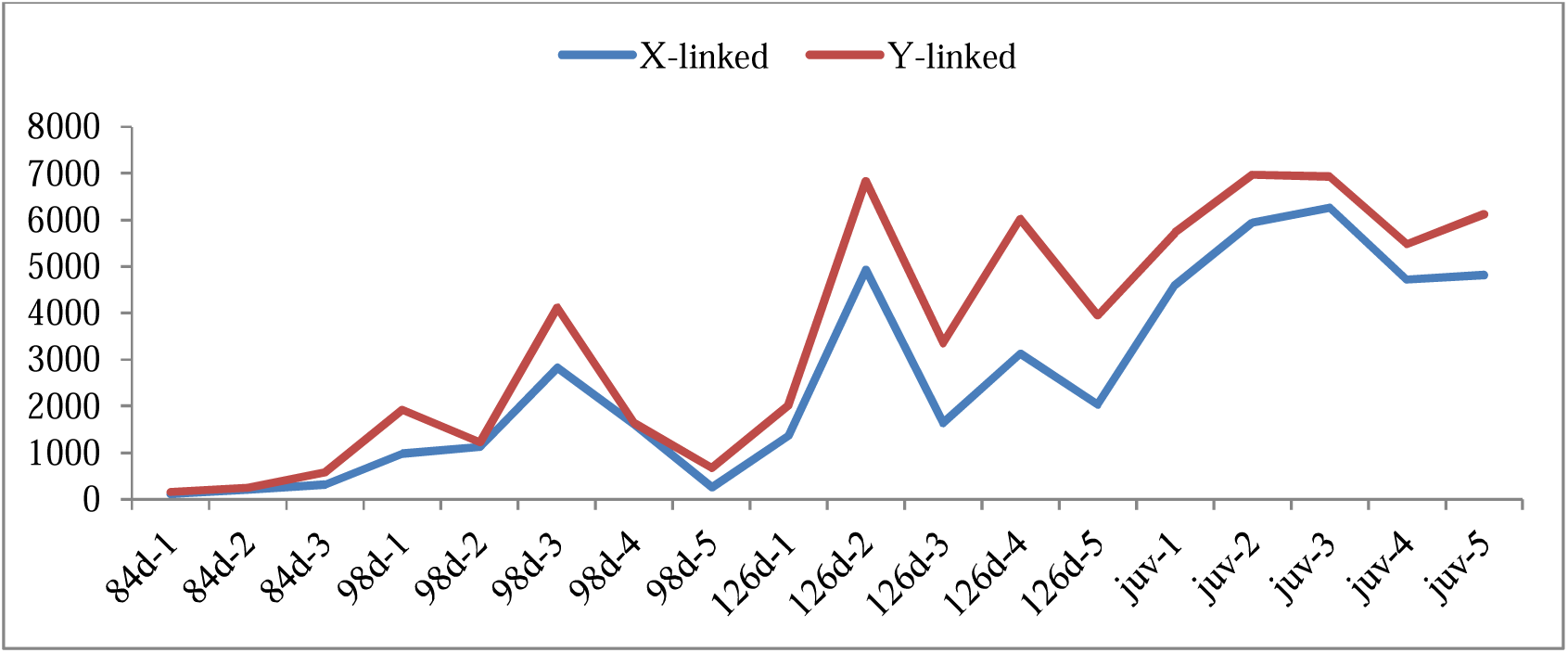
Number of counts for the X-linked and Y-linked allelic variants of the *fshr* gene across different gonad developmental stages in *Solea senegalensis*.

Additionally, marker genes of germinal cell proliferation (*gsdf* and *vasa*) and of male (*sox9a* and *amh*) and female (*cyp19a1a*) gonadal differentiation, were evaluated by qPCR. While adult male and females hardly showed differential expression for the genes evaluated, juveniles showed the greatest differentiation for most of them, excluding *cyp19a1a* and *vasa* (maximum at 126 dpf). Remarkably, *sox9a* showed much higher expression in male juveniles, which contrasted with its higher expression in females from previous stages. A progressive increase in expression was observed for *cyp19a1a* and *vasa* from 84 dpf to 126 dpf, while *amh* expression peaked at 98 dpf and *gsdf* at 84d. Germinal cell proliferation gene markers showed opposite patterns at the initial stages, *vasa* was always overexpressed in females (significantly at 98 and 126 dpf), while *gsdf* was overexpressed in males at 86 and 98 dpf. Sex markers showed the expected patterns for *amh*, overexpressed in males at 86 and 98 dpf, and *cyp19a1a*, overexpressed in females at all stages since 86 dpf; however, *sox9a* was overexpressed in females, suffering an abrupt change in juveniles, as outlined before.

## Discussion

### A new Senegalese sole genome assembly

The new genome assembly of the Senegalese sole is amongst the most contiguous fish genomes assembled to date (Ramos & Antunes 2022) and meets the standards outlined by the EBP initiative (https://www.earthbiogenome.org/assembly-standards). The contiguity achieved was facilitated by the small proportion of low complexity sequences; repetitive elements only constitute 8.0% of the *S. senegalensis* genome, a similar proportion to that reported in other flatfish species (Chen et al., 2014; Figueras et al., 2016). The new *S. senegalensis* assembly comprises 21 scaffolds corresponding to the n = 21 chromosomes of its haploid karyotype (Vega et al., 2002) and contains 99.02% of the whole assembly, higher than other flatfish genomes recently reported (Einfeldt et al., 2021; Lü et al., 2021; Martínez et al., 2021; Ferchaud et al., 2022). These 21 scaffolds were associated to the corresponding chromosomes of this species using a BAC-FISH approach (average 6.6 BACs / chromosome), thus providing a sound reference for comparative genomics and for studying the genetic architecture of relevant traits, such as sex determination. The new assembly displayed a major one to one chromosome correspondence with the recently reported genome by Guerrero-Cozar et al. (2021), and most of the discordances were related to orientation of minor scaffolds due to the higher fragmentation and less contiguity of the previous assembly (∼1937 scaffolds; 90 % of the assembly integrated in chromosomes). This upgraded Senegalese sole genome allowed the identification and annotation of 24,264 protein-coding genes encoding for 37,259 unique protein products (85% with functional annotation), along with 52,888 non-coding RNAs (6,871 long non-coding and 46,017 small RNAs).

### Comparative genomics within Pleuronectiformes: orthology

Orthology relationships were established across the six species of the order with chromosome-level assemblies to understand the chromosome reorganizations in the evolutionary history of Senegalese sole. As previously reported for Acanthoptherygii (Maroso et al., 2018), a general macrosyntenic conservation across flatfish species was confirmed. The pair-wise collinearity between genomes was similar to that reported by Lü et al. (2021) and related to their phylogenetic distance. Four consistent reorganization patterns in the evolution of the Senegalese sole genome / karyotype were identified: i) centric fusions of two acrocentric chromosomes giving rise to metacentric chromosomes, either lineage-specific (Sse C1, Sse C4) or representing independent events in different lineages (Sse C2 *vs* Sma C2); ii) pericentric inversions of acrocentric chromosomes originating biarmed chromosomes (C3, C5-C9); iii) translocations involving small or medium chromosome fragments (e.g. intercalary syntenic blocks of Sse C16 spread over *S. maximus* Sma C16 comprehending 18.5% of the chromosome markers); and iv) extensive intrachromosomic shuffling of microsyntenic blocks, as reported in this order (Maroso et al., 2018; Guerrero-Cózar et al. 2021). The robust mapping integration and comprehensive proteome obtained confirmed previous observations and provided new insights on the chromosome evolutionary trends in the order (Robledo et al., 2017; Lü et al., 2021).

### The master SD gene of Solea senegalensis

Guerrero-Cozar et al. (2021) reported a broad genomic region (1.4 Mb) associated with sex at SseLG18 of *S. senegalensis* (Sse C12 in our study) using five full-sib families, compatible with a nascent XX / XY system. They suggested a SD mechanism with incomplete penetrance influenced by environmental factors since sex association was incomplete, and proposed *fshr* as the most promising master SD gene considering its role in gonad development and previous data from other species (Curzon et al., 2021). Here, we demonstrated the presence of a Y-linked *fshr* allelic variant (*fshry*) in the Senegalese sole, compatible with the proposed XX / XY SD system and congruent with phenotypic sex in 98% of the individuals analysed. Genetic differentiation between males and females was essentially restricted to the *fshr* gene (∼9 kb). The high number of diagnostic variants, heterozygous in males and homozygous in females, detected in the *fshry* coding region (33), 24 of them representing non-synonymous substitutions, suggested that it could represent a non-functional allele. Nonetheless, *fshry* was not only expressed at nearly all stages of gonadal development, but its expression was always higher than that of the X-linked allele (*fshrx*). This hormone receptor is a G protein-coupled with seven-transmembrane domains linked to an adenyl cyclase for intracellular transduction of the signal (Levavi-Sivan et al., 2010). Interestingly, non-synonymous variants were mostly concentrated at exon 14 and to an extent at exon 1, which are part of its intracellular and extracellular domains, respectively, related to signal transduction and hormone reception. Moreover, we could not detect evidence of a duplication of the *fshr* gene in the Senegalese sole male genome reported by Guerrero-Cozar et al. (2021), and therefore, the *fshry* allele appears to be the key factor for the male fate of the undifferentiated primordium. A similar SD mechanism has been reported in gray mullet, where an *fshry* allele including only two non-synonymous variants was identified as responsible for the male fate of the undifferentiated primordium (Curzon et al., 2021). It can be speculated that the mechanism driven by the *fshry* allele might be at a more advanced evolutionary stage in Senegalese sole, considering the 24 non-synonymous variants detected, and in fact, Ferrareso et al. (2021) has reported an interpopulation variation of the *fshry* in gray mullet, suggesting an incomplete penetrance and a recent evolutionary origin of the SD gene.

The expression of *fshr* measured by both qPCR and RNAseq was consistently higher in males than in females from 84 dpf (undifferentiated gonads), and this difference increased progressively until the juvenile stage, which displayed the largest difference. In females, expression was negligible at most stages, with the exception of 98 dpf and 126 dpf, and in adults expression was nearly undetectable in both males and females. The higher expression of *fshr* in males has also been reported in other fish species like clownfish (Kobayashi et al., 2017) and gray mullet (Dor et al., 2020). However, deleterious mutations or knock-out of the *fshr* gene have been shown to affect ovary development by impairing folliculogenesis in medaka and other vertebrates (Zariñán et al., 2010), and have also been connected with female to male sex reversal in zebrafish and medaka (Murozumi et al., 2014; Zhang et al., 2015; Chu et al., 2015). Furthermore, *fshr* has been suggested as the masculinisation transducer of cortisol via suppression of germinal cell proliferation in response to high temperature during the sex determinant period (Hayashi et al., 2010).

The analysis of *fshr* and other key genes at the initial stages of gonad differentiation in Senegalese sole revealed a progressive upregulation of *fshr, amh* and *gsdf* in males, whereas *cyp19a1* and *vasa* displayed a similar upregulation pattern in females. It should be noted that the expression of *amh*, a maker gene of testis development, also reported as a master SD gene in several fish species (or its receptor *amhr*), is down-regulated by the follicle stimulating hormone (Sambroni et al., 2013).

All in all, our data strongly support *fshr* as the master SD gene of Senegalese sole and suggest that the *fshry* allele could be hampering the action of the follicle stimulating hormone (FSH), driving the undifferentiated gonad toward testis, potentially by avoiding the suppression of *amh* activity. Although we cannot exclude the existence of alternative splicing variants of this gene triggering the testis pathway, our data showed correlated expression profiles across the 14 exons in males and females, and between the *fshrx* and *fshry* variants in males, which does not support such mechanism. We hypothesize that the presence of 11 non-synonymous variants in the intracellular part of the receptor should affect the transduction signal of *fshr*. Furthermore, the higher expression of the *fshry* allele, along with the non-synonymous variants occurring in the extracellular hormone-reception domain of the protein (exon 1), could increase its affinity for FSH, thus sequestering molecules and impairing its action. The study of 3D models of the *fshr* variants and their interaction with the hormone and with the internal transducer could aid to confirm this hypothesis

### Diversification of the SD gene in Pleuronectiformes

Based on synteny data, the Sse C12 of Senegalese sole (XX/XY system) showed no orthology to other reported SD chromosomes across flatfish species, irrespective of their chromosome systems, which gives additional support to its novelty within Pleuronectiformes. As reported in turbot (Martínez et al. 2021), the low differentiated Z and W chromosomes of tongue sole (Chen et al. 2014) matched to a single Senegalese sole chromosome, Sse C6, which is syntenic to the SD-Hst9 of the Pacific halibut (Drinan et al. 2018). This reinforces the orthology of sex chromosomes of tongue sole and Pacific halibut (Martínez et al. 2021), not conserved in other flatfish (Martínez et al. 2021). Interestingly, Sse C12 is orthologous to the turbot Sma C18, where a minor SD-QTL was identified and where sex associated markers were detected in the brill (*S. rhombus*), a congeneric XX/XY species within Scopthalmidae (Taboada et al. 2014). Moreover, Sse C12 is syntenic to the chromosome 21 of Greenland halibut (*Reinhardtius hippoglossoides*), where *sox9a* was suggested as a possible SD candidate gene along with *gdf6* and *sox2* in the chromosome 10 of this XX/XY species (Ferchaud et al. 2022). This latter gene would point to a similar SD system as reported in the turbot (Martínez et al. 2021). The syntenic relationships between Senegalese sole and other flatfish adds evidence of the huge heterogeneity of SD systems in

Pleuronectiformes, even between closely related species (Drinan et al. 2018; Einfeldt et al. 2021; Martínez et al. 2021; Ferchaud et al. 2022), but also highlights the recruitment of major SD drivers from common gene families (e.g., *fshr, dmrt, amh* or *sox*) across different fish and vertebrate species (Ferraresso et al. 2021; Martínez et al. 2021; Ferchaud et al. 2022).

## Conclusions

The chromosome-level Senegalese sole genome assembly reported in this study is among the most contiguous fish assemblies to date. Its integration with previous resources has generated a robust genomic framework for future studies in this emerging aquaculture species. Comparative genomics including other chromosome-level flatfish assemblies confirmed the macrosyntenic pattern in the order and identified the main chromosome evolutionary trends that configured the karyotype of Senegalese sole. This new high-quality assembly enabled the identification of the master SD gene of the species, the follicle stimulating hormone receptor (*fshr*), a new SD gene reported for the first time in Pleuronectiformes, which has also been recently suggested as the master SD gene in gray mullet. The soundest hypothesis is that the Y-linked *fshry* variant might be hampering the function of the FSH hormone, impeding the down-regulation of *amh* and driving the undifferentiated gonad toward testis. Last but not least, an important outcome of this study is the validation of a practical tool for sexing, which would facilitate the production of all-female populations for industry.

## Acknowledgements

This study was supported by the Spanish Ministry of Economy and Competitiveness, FEDER Grants (RTI2018-097110-B-C21 and RTI2018-096847-B-C22), Junta de Andalucía-FEDER Grants (RTI2018-096847-B-C21 and P20-00938) and the European Union’s Horizon 2020 research and innovation programme under grant agreement No 81792 (AQUA-FAANG). We thank Geneaqua SL for their participation and financial support on sequencing. We acknowledge the bioinformatic support of the Centro de Supercomputación de Galicia (CESGA).

## Supplementary information

### Supplementary Tables

**Table S1:** Genome assembly statistics of *Solea senegalensis*; fSolSen1_LG: set of contigs anchored to linkage groups (LG) of the genetic map.

**Table S2:** Statistics of TE-derived sequence and other simple repeats in the genome of *Solea senegalensis*.

**Table S3:** Genome annotation statistics of protein coding genes in the genome of *Solea senegalensis*

**Table S4:** Absolute number of reads and averages obtained by the 2b-RAD method for SNP genotyping in three families (F1, F2, F3) of *Solea senegalensis* and their offspring across the different filtering steps of the established pipeline from the initial raw reads to the final valid alignment for constructing a highly dense genetic map.

**Table S5:** Statistics of the genetic maps constructed in *Solea senegalensis* using three full-sib families. Maps were constructed via male and via female in each family and the consensus per sex and for the whole species obtained. The correspondence between linkage groups in the consensus map and that reported by Guerrero-Cózar et al. (2021) is also provided. LG codes were arranged from the longest to the shortest within each map, but their correspondence was established according to the codes of the consensus map, which in turn followed the chromosome number of the karyotype after mapping integration. LG: linkage group; families: F1, F2 and F3; male: M; female: F.

**Table S6:** Marker positions for all genetic maps constructed in *Solea senegalensis* and their integration by sex and species.

**Table S7:** List of anchored scaffolds of the *Solea senegalensis* genome on the genetic map indicating the orientation and size.

**Table S8:** Comparative statistics of the *Solea senegalensis* genome with other pleuronectiform chromosome-level genomes.

**Table S9:** One hundred and forty-one BAC clones used for integrating cytogenetic, genetic and physical maps in *Solea senegalensis*. Notice that some clones matched to several regions, either in different or the same chromosome, among them, the 5S rDNA gene clusters

**Table S10:** Orthology statistics for: A) six commercial flatfish proteomes with genomes resolved at chromosome-level; and B) seven fish species, which include the six flatfish and the model teleost *Danio rerio* (Bioproject PRJNA11776). Flatfish species studied: *Solea senegalensis* (Soleidae), *Cynoglossus semilaevis* (Cynoglossidae), *Hippoglossus hippoglossus* and *H. stenolepis* (Pleuronectidae), *Paralichthys olivaceus* (Paralychthydae) and *Scophthalmus maximus* (Scophthalmidae). Proteome data came from this study and PRJNA251742, PRJNA562001, PRJNA622249, PRJNA73673 and PRJNA631898 bioprojects, respectively.

**Table S11A:** Single-copy orthologous genes anchored to chromosomes in the six flatfish species studied, including the functional annotation in the *Solea senegalensis* genome.

**Table S11B:** Single-copy orthologous genes in the six flatfish species studied and the model teleost *Danio rerio*.

**Table S12**. Chromosome relationships among the six chromosome-level flatfish genomes, based on syntenic single-copy orthologous genes (Table S11A). The color and chromosome identity for each species from Figure 3. Sse_*Solea senegalensis* (Family Soleidae), Cse_*Cynoglossus semilaevis* (Cynoglossidae), Sma_*Scophthalmus maximus* (Scophthalmidae), Pol_*Paralichthys olivaceus* (Paralychthydae), Hst_*Hippoglossus stenolepis* (Pleuronectidae), Hhi_*Hippoglossus hippoglossus* (Pleuronectidae).

**Table S13:** Wright F-statistics for male and female populations per SNP and using 50 SNP-sliding windows averaged over F_IS_ and F_ST_ across the contig 19 anchored to C12 of the *Solea senegalensis* genome, where the *fshr* gene is located

**Table S14:** Characteristics of SNPs localized in the follicle stimulating hormone receptor (fshr) gene in six males and six females resequenced at 20x coverage using the *Solea senegalensis* assembled genome; diagnostic SNPs are homozygous in females and heterozygous in males consistent with a XX / XY sex determining system; in the last row it is indicated if SNPs determine aminoacid substitutions (non-synonymous) or not (synonymous); the boundaries of exons are highlighted in bold type.

**Table S15:** Sets of primers designed with Primer 3 to develop a molecular tool for sexing in *Solea senegalensis* using diagnostic markers between males (heterozygous) and females (homozygous) located at exon 14 of the *fshr* gene.

**Table S16:** Genotypes and allelic counts of diagnostic SNPs located in the *fshr* gene (exons, 5’ and 3’ UTR, introns; detailed information in Table S14) from gonad RNAseq data of five males (M) and five females (F) sampled across gonad development of *Solea senegalensis*, from the initial undifferentiated or low differentiated stages (84D, 98D and 126D post fertilization) until juveniles and adults; SNP ID makes reference to the contig and position where SNPs are located in the genome; REF and ALT alleles refers to the allele in the genome (0) and the alternative allele (1) detected after resequencing six females and six males; Genotypes: homozygous in females (0/0 or 1/1) and heterozygous in males (0/1); ./. missing genotypes because allelic counts did not reach the minimum threshold (8 reads); allelic counts: the first number refers to the “0” allele and the second, separated by semicolon, to “1” allele; Colours: pink (missing genotypes), green (valid genotypes for counting), pink (females); blue (males); red (individuals not considered because they did not reach a minimum genotyping data).

### Supplementary Figures

**Fig. S1:** K-mer distribution on: A) initial Illumina reads; B) final assembly

**Fig. S2:** LASTZ plots between scaffolds / chromosomes (this study; ordinates) and pseudo-chromosomes (Guerrero-Cózar et al., 2021; abscisse) of *Solea senegalensis* genome assemblies; A) psedochromosomes 1-12; B) pseudochromosomes 13-21.

**Fig. S3A:** Results of double (a – e) and multicolor FISH (f – h) with *Solea senegalensis* BACs:. (a) 10L10 (red) / 67N4 (green) for chromosome 1 (C1), (b) 7H22 (red) / 47G8 (green) for C7, (c) 15I19 (red) / 57N7 (green) for C10, (d) 71N11 (green) / 54E18 (red) for C16, (e) 55B12 (red) / 72O12 (green) for C21, (f) 42D4 (blue) / 65E23 (red) / 21I14 (green) for C2 and 54E18 (pink) located in C16, (g) 36H3 (pink) / 67K3 (blue) / 39G22 (green) located in C4 and 65E23 (red) for C2, and (h) 38H3 (green) / 62G15 (blue) / 54G7 (red) positioned in C19 and 67N4 (pink) for C1. Bar = 2 μm (referred to large metacentric C1).

**Fig. S3B:** Representation of the *Solea senegalensis* chromosomes with BACs used for mapping integration.

**Fig. S4**. Phylogenetic trees for orthogroups present in six flatfish species inferred using OrthoFinder (Emms and Kelly, 2019). A) Example of a many-to-many gene tree; B) Example of one-to-one (single copy orthologous) gene tree; C) Unrooted species tree using single copy orthologous genes in six flatfish species: *Solea senegalensis* (Soleidae), *Cynoglossus semilaevis* (Cynoglossidae), *Hippoglossus hippoglossus* and *H. stenolepis* (Pleuronectidae), *Paralichthys olivaceus* (Paralychthydae) and *Scophthalmus maximus* (Scophthalmidae); D) Rooted species tree based on single-copy conserved orthologs in all flatfish species and *Danio rerio*.

**Fig. S5**. Pairwise circos representations of orthologous gene relationships between *Solea senegalensis* (Soleidae) chromosomes (right) and each of the five chromosome-level flatfish genomes studied (left): *Scophthalmus maximus* (Scophthalmidae), *Cynoglossus semilaevis* (Cynoglossidae), *Paralichthys olivaceus* (Paralychthydae), *Hippoglossus hippoglossus* and *H. stenolepis* (Pleuronectidae).

**Figure S6**. Syntenic relationships based on orthologous genes between the metacentric chromosome 2 of *Solea senegalensis* (Sse C2; y-axis) and the metacentric chromosome 2 of turbot (Sma2; x-axis), according to the consistent syntenies with orthologous acrocentric chromosomes in other flatfish: *Paralichthys olivaceus* (Pol9), *Hippoglossus stenolepis* (Hst23*), H. hippoglossus* (Hhi18) and *Cynoglossus semilaevis* (Cse1); B) Pol16-Hst22-Hi23-Cse8.

**Figure S7**. Pairwise gene orthology plots between the *Solea senegalensis* metacentric chromosome Sse C3 (Y-axis) and acrocentric chromosomes of the other five flafish species (X-axis). Chromosome positions are given in base pairs; A) Sse3 *vs. Cynoglossus semilaevis* (Cse 1); B) Sse3 *vs. Scophthalmus maximus* (Sma17); C) Sse3 *vs. Paralychthys olivaceous* (Pol8); D) Sse3 *vs. Hippoglossus hippoglossus* (Hhi22); E) Sse3 *vs. H. stenolepsis* (Hst10).

**Figure S8**. Syntenic relationships between *Cynoglossus semilaevis* C5 (Cse5; x-axis) and other flatfish species (y-axis) showing extensive reorganizations at microsyntenic scale.

**Figure S9**. Macroscopic anatomy and topography of *Solea senegalensis* gonads of individuals at 128, 98 and 86 dpf. (A, B) 128 dpf female sole; (C) 98 dpf individual - unidentified sex; (D, E) 128 dpf presumably sole male identified by ‘lacking a female gonad’. (F) 96 dpf individual with unidentified sex. Insets: site of gonad. Scale bars: 600 μm (A); 400 μm (B-F).

**Figure S10**. Histological sections of adult and juvenile *Solea senegalensis* gonads. (A-C) Adult female. (D-F) Juvenile female. (G-I) Adult male. (J-L) Juvenile male. (A-F) (*) Atresia stages; (^) Post-ovulatory follicles; Arrow: Nuclei; Arrowhead: Nucleolus; (1) Oogonia; (2) Early oocyte; (3) Late oocyte. (G-L) White arrows: Show the radial disposition of seminiferous lobules (*) from the central medulla (m) to the cortex (c) and tunica albuginea (ta); (1) and black arrowhead: Spermatocysts; 2 and black arrow: Spermatozoa; Black square: Intersticial tissue. Stain: Hematoxilin-Eosin (HE). Scale bars: 250 μm (A,D,G); 100 μm (B,C,E,H,J,K); 50 μm (F,I,L).

**Figure S11**. Histological sections of *Solea senegalensis* female gonads of individuals at 126, 98 and 86 dpf. (A) In 128 dpf female sole, previtellogenic oocytes can be visualized; (B,C) Gonad of 98 and 86 dpf females, respectively, both undifferentiated, although at 98 dpf there are more potential-oocyte-cells. (D-F) Higher magnification of A-C, respectively. Gonads of 98 and 86 dpf individuals correspond to females identified with the SS-sex. Stain: HE. Scale bars: 250 μm (A); 100 μm (B); 50 μm (C,D,E,F).

**Figure S12:** Histological sections of *Solea senegalensis* male gonads of individuals at 126, 98 and 86 dpf. (A) 128 dpf; (B) 98 dpf; (C) 86 dpf; (D, E) Higher magnification of A, B, respectively. The three stages are undifferentiated, with the oldest one (126 dpf) showing more potentially-spermatogonia cells. All samples were genotyped with the SS-sex marker. * Gonad; K: kidney; c: Cartilage. Stain: HE. Scale bars: 100 μm (A, B, C); 50 μm (D, E).

**Figure S13:** RNAseq data of the *fshr* gene in *Solea senegalensis* males and females across different gonad developmental stages (abscissas) using a count log scale (ordinates).

**Figure S14:** Read counting profiles across the 14 exons of the follicle stimulating hormone receptor gene in males and females at 98 dpf, 126 dpf and juveniles (only males).

