## Supplementary material for "A chromosome-level genome assembly enables the identification of the follicle stimulating hormone receptor as the master sex determining gene in *Solea senegalensis*": Table S2

**Table S2**. Statistics of TE-derived sequence and other simple repeats in the genome of *Solea senegalensis*.

| Class | Order | *Clade/Superfamily* | No. fragments | Total length (kbp) | Genome fraction  (%) |
| --- | --- | --- | --- | --- | --- |
| RTs |  |  | 76621 | 12136 | 2.29 |
|  | DIRS |  | 2231 | 274 | 0.05 |
|  |  | Ngaro | 1344 | 154 | 0.03 |
|  |  | DIRS | 887 | 120 | 0.02 |
|  | LINE |  | 28674 | 6436 | 1.21 |
|  |  | L2 | 14588 | 3055 | 0.58 |
|  |  | L1 | 5290 | 1051 | 0.20 |
|  |  | RTE | 3801 | 908 | 0.17 |
|  | LTR |  | 28206 | 3291 | 0.62 |
|  |  | Gypsy | 9027 | 1781 | 0.34 |
|  |  | ERV1 | 8540 | 690 | 0.13 |
|  |  | ERVK | 6018 | 389 | 0.07 |
|  |  | Pao | 1283 | 256 | 0.05 |
|  |  | Copia | 96 | 20 | 0.00 |
|  |  | ERVL | 332 | 19 | 0.00 |
|  |  | ERV4 | 200 | 12 | 0.00 |
|  |  | Ginger | 32 | 5 | 0.00 |
|  |  | ERVL-MaLR | 32 | 2 | 0.00 |
|  | PLE | Penelope | 1613 | 183 | 0.03 |
|  | SINE |  | 15897 | 1952 | 0.01 |
|  |  | tRNA-V | 6438 | 1175 | 0.22 |
|  |  | MIR | 3990 | 458 | 0.09 |
|  |  | tRNA-Core | 3384 | 441 | 0.08 |
|  |  | Mermaid | 642 | 55 | 0.01 |
|  |  | L2 | 875 | 50 | 0.01 |
|  |  | tRNA | 606 | 45 | 0.01 |
|  |  | tRNA-Core-L2 | 606 | 45 | 0.01 |
|  |  | tRNA-V-CR1 | 373 | 37 | 0.01 |
|  |  | 5S-Deu-L2 | 358 | 23 | 0.00 |
|  |  | 5S-Sauria-RTE | 135 | 15 | 0.00 |
|  |  | tRNA-V-Core-L2 | 122 | 13 | 0.00 |
|  |  | tRNA-L2 | 148 | 8 | 0.00 |
|  |  | tRNA-C | 126 | 7 | 0.00 |
|  |  | B4 | 63 | 4 | 0.00 |
|  |  | ID | 43 | 2 | 0.00 |
|  |  | tRNA-Deu-L2 | 35 | 2 | 0.00 |
|  |  | 7SL | 7 | 1 | 0.00 |
|  |  | tRNA-RTE | 22 | 1 | 0.00 |
| DNA |  |  | N.A. | 13557 | 2.56 |
|  | Helitron | Helitron | 5573 | 513 | 0.10 |
|  | TIR |  | 60743 | 13041 | 2.12 |
|  |  | hAT | 31015 | 2473 | 0.47 |
|  |  | Tc1-Mariner | 10861 | 1931 | 0.36 |
|  |  | EnSpm | 7360 | 605 | 0.11 |
|  |  | Maverick | 4183 | 371 | 0.07 |
|  |  | PIF-Harbinger | 1875 | 208 | 0.04 |
|  |  | Kolobok-T2 | 2213 | 156 | 0.03 |
|  |  | Dada | 2160 | 156 | 0.03 |
|  |  | PiggyBac | 825 | 68 | 0.01 |
|  |  | Academ | 113 | 17 | 0.00 |
|  |  | Harbinger | 80 | 8 | 0.00 |
|  |  | PIF-ISL2EU | 23 | 2 | 0.00 |
|  |  | MITEs | N.A. | 7043 | 1.33 |
|  | Unclassified |  | 9102 | 793 | 0.15 |
| Total interspersed repeats | |  | N.A. | 26486 | 5.00 |
|  | Other motifs | Small RNAs | 2065 | 156 | 0.03 |
|  |  | Satellites | 5432 | 664 | 0.13 |
|  |  | Simple repeats | 358944 | 15954 | 3.01 |
|  |  | Low complexity | 33492 | 1860 | 0.35 |

Note. N.A.- Not available. Superfamilies contributing < 1 kb of genomic sequence were not included.
