## Supplementary material for "A chromosome-level genome assembly enables the identification of the follicle stimulating hormone receptor as the master sex determining gene in *Solea senegalensis*": Table S7

**Table S7**. List of anchored scaffolds of the *Solea senegalensis* genome on the genetic map indicating the orientation and size.

| **LG / chrom** | **pseudo-chromosomes** | **Scaffold** | **Orientation** | **size (bp)** | **size /chrom** |
| --- | --- | --- | --- | --- | --- |
| 1 | 1 | SolSen1_s38c1 | Reverse | 781045 |  |
|  |  | SolSen1_s13c1 | Reverse | 23439045 |  |
|  |  | SolSen1_s15c1 | Forward | 22142923 | 46363013 |
| 2 | 3 | SolSen1_s23c1 | Reverse | 13730948 |  |
|  |  | SolSen1_s30c1 | Reverse | 3794735 |  |
|  |  | SolSen1_s20c1 | Reverse | 18999240 | 36524923 |
| 3 | 16 | SolSen1_s10c1 | Reverse | 24069211 |  |
|  |  | SolSen1_s56c1 | Forward | 286902 | 24356113 |
| 4 | 2 | SolSen1_s46c1 | Forward | 439503 |  |
|  |  | SolSen1_s07c1 | Reverse | 24577862 |  |
|  |  | SolSen1_s22c1 | Reverse | 14663850 | 39681215 |
| 5 | 21 | SolSen1_s14c1 | Reverse | 22452756 | 22452756 |
| 6 | 5 | SolSen1_s42c1 | Forward | 522248 |  |
|  |  | SolSen1_s01c1b | Reverse | 21630749 |  |
|  |  | SolSen1_s28c1 | Reverse | 8058068 | 30211065 |
| 7 | 7 | SolSen1_s09c1 | Reverse | 24206605 |  |
|  |  | SolSen1_s43c1 | Reverse | 507339 |  |
|  |  | SolSen1_s27c1 | Reverse | 8362269 | 33076213 |
| 8 | 10 | SolSen1_s04c1 | Reverse | 28132043 | 28132043 |
| 9 | 12 | SolSen1_s45c1 | Reverse | 446947 |  |
|  |  | SolSen1_s29c1 | Forward | 5975391 |  |
|  |  | SolSen1_s01c1a | Reverse | 12020340 |  |
|  |  | SolSen1_s25c1 | Reverse | 8875931 |  |
|  |  | SolSen1_s41c1 | Forward | 569458 | 27888067 |
| 10 | 20 | SolSen1_s37c1 | Forward | 856747 |  |
|  |  | SolSen1_s48c1 | Forward | 395635 |  |
|  |  | SolSen1_s16c1 | Reverse | 20937294 |  |
|  |  | SolSen1_s47c1 | Forward | 414049 |  |
|  |  | SolSen1_s50c1 | Forward | 327553 | 22931278 |
| 11 | 17 | SolSen1_s12c1 | Forward | 23564453 | 23564453 |
| 12 | 18 | SolSen1_s33c1 | Reverse | 2416913 |  |
|  |  | SolSen1_s19c1 | Reverse | 19354633 | 21771546 |
| 13 | 11 | SolSen1_s18c1 | Forward | 19885849 |  |
|  |  | SolSen1_s26c1 | Reverse | 8756652 | 28642501 |
| 14 | 6 | SolSen1_s52c1 | Forward | 317263 |  |
|  |  | SolSen1_s64c1 | Forward | 224809 |  |
|  |  | SolSen1_s24c1 | Reverse | 13270792 |  |
|  |  | SolSen1_s21c1 | Reverse | 15211903 | 29024767 |
| 15 | 19 | SolSen1_s32c1 | Reverse | 3259233 |  |
|  |  | SolSen1_s17c1 | Reverse | 20639214 |  |
|  |  | SolSen1_s36c1 | Forward | 889564 | 24788011 |
| 16 | 9 | SolSen1_s05c1 | Reverse | 27800896 | 27800896 |
| 17 | 13 | SolSen1_s63c1 | Forward | 229283 |  |
|  |  | SolSen1_s11c1 | Forward | 23966579 | 24195862 |
| 18 | 15 | SolSen1_s08c1 | Reverse | 24552006 | 24552006 |
| 19 | 4 | SolSen1_s35c1 | Reverse | 1032886 |  |
|  |  | SolSen1_s02c1 | Reverse | 30103958 | 31136844 |
| 20 | 8 | SolSen1_s34c1 | Forward | 1773698 |  |
|  |  | SolSen1_s03c1 | Reverse | 28862912 | 30636610 |
| 21 | 14 | SolSen1_s06c1 | Forward | 25475828 |  |
|  |  | SolSen1_s31c1 | Forward | 3674203 | 29150031 |
| **Total** |  |  |  | 606880213 |  |
