## Supplementary material for "A chromosome-level genome assembly enables the identification of the follicle stimulating hormone receptor as the master sex determining gene in *Solea senegalensis*": Table S8

| **Table S8.** Comparative statistics of the *Solea* *senegalensis* genome with other pleuronectiform chromosome-level genomes. | | | | | | | | | | |
| --- | --- | --- | --- | --- | --- | --- | --- | --- | --- | --- |
|  | ***Solea senegalensis*** | | ***Platychthys stellatus*** | ***Scophthalmus maximus*** | ***Cynoglossus semilaevis*** | ***Hippoglossus hippoglossus*** | ***Hippoglossus stenolepis*** | ***Reinhardtius hippoglossoides*** | ***Verasper variegatus*** | ***Paralichthys olivaceous*** |
|  | **this study** | ***Guerrero-Cózar et al.* (2021)** | Lü et al (2021) | Martínez et al. (2021) | Chen et al (2014) | Einfeldt et al. (2021) | NCBI Bioproject | Ferchaud et al., 2022 | Zhao et al., 2021 | Shao et al., 2017 |
| Chromosome number | 21 | 21 | 24 | 22 | 21 | 24 | 24 | 24 | 23 | 24 |
| Bioproject |  | PRJNA643826 | PRJNA592732 | PRJNA631898 | PRJNA251742 | PRJNA562001 | PRJNA622249 | PRJNA499109 | PRJNA634516 | PRJNA73673 |
| GC_content (%) | 40.90 | 40.92 | 42.66 | 43.38 | 41.27 | 42.27 | 42.24 | 42.65 | 42.11 | 41.62 |
| Contig / scaffold N50 (Mb) | 28.6 | 26 | 25.1 | 25.94 | 0.5 | 26.31 | 24.98 | 25.01 | 24.76 | 3.81 |
| Largest contig / scaffold (Mb) | 36.20 | 42.92 | 31.75 | 32.11 | NA | 31.67 | 32.41 | 31.98 | 31.93 | NA |
| Total length (Mb) | 613.0 | 603.53 | 609.99 | 556.7 | 470.19 | 596.79 | 594.14 | 598.52 | 545.34 | 545.77 |
| % assembled in chromosomes | 99% | 90% | 92% | 96% | 94% | 94% | 98% | 96% | 99% | 98% |
| Contig / scaffold number | 81 | 1,937 | 31,621 | 127 | 31,181 | 57 | 120 | 1590 | 57 | 7,202 |
