## Supplementary figures and images for "A chromosome-level genome assembly enables the identification of the follicle stimulating hormone receptor as the master sex determining gene in *Solea senegalensis*"

### Fig. S2A, S2B

#### Slide 1
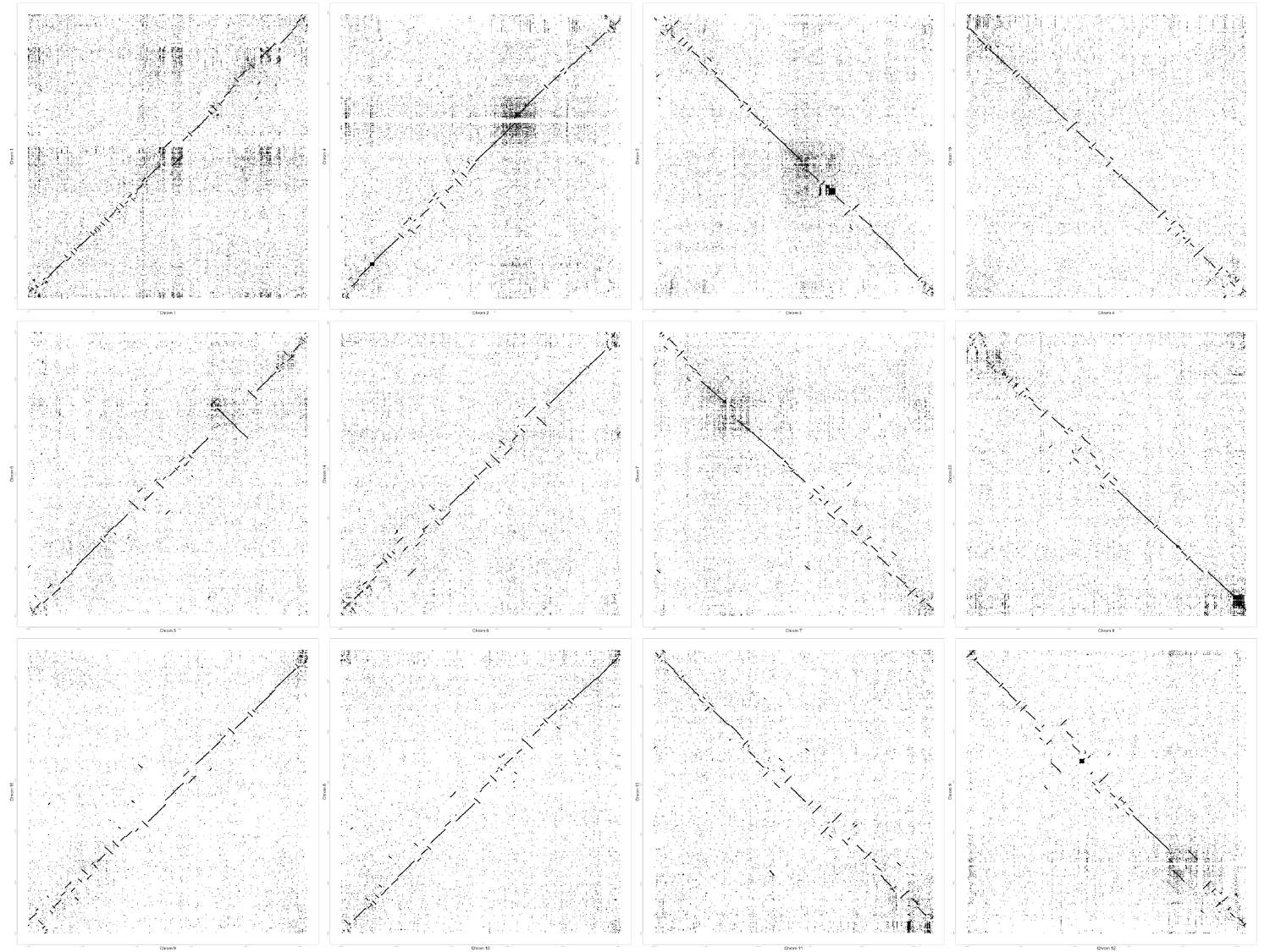

#### Slide 2
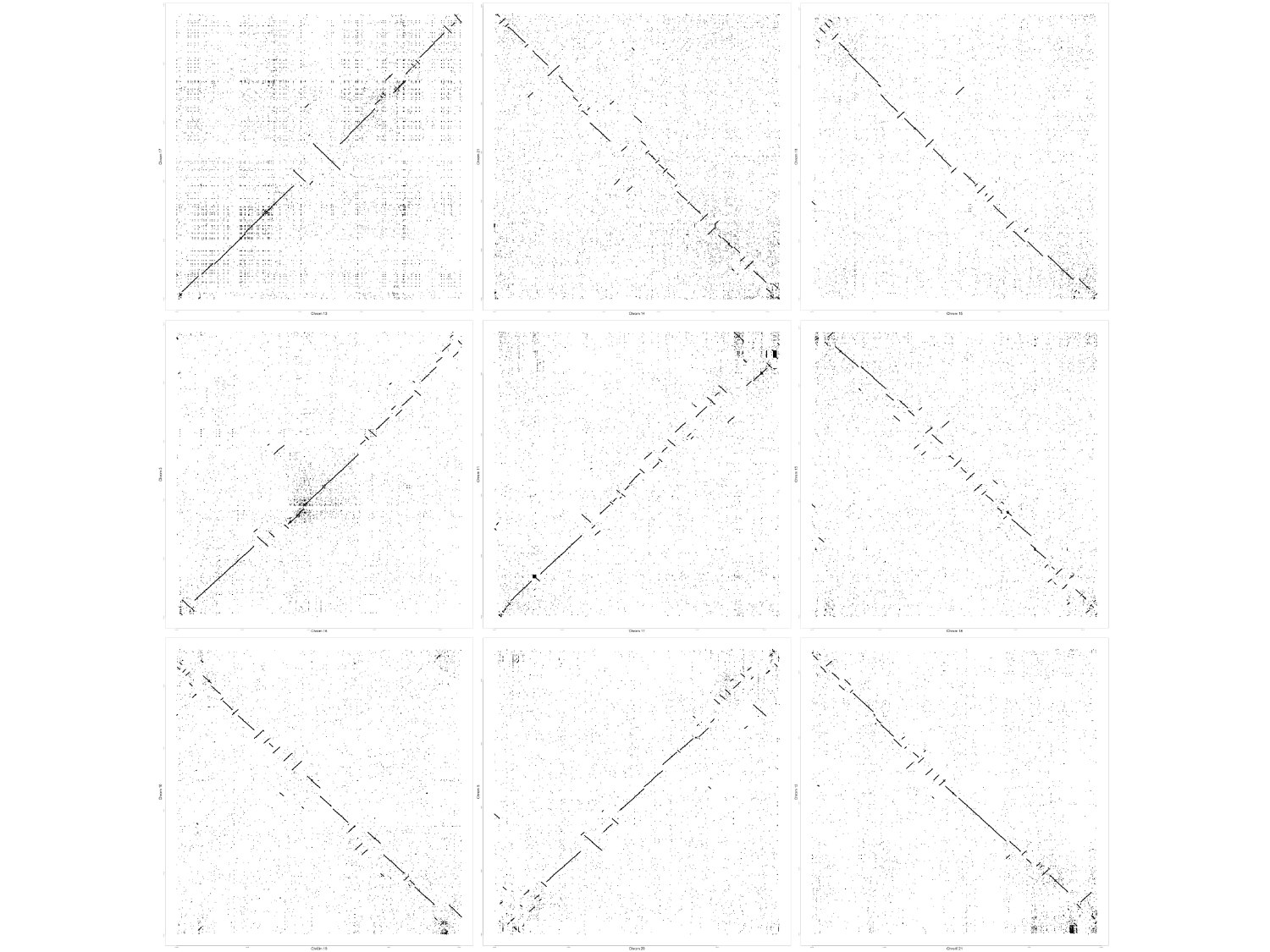

### Fig. S3B

#### Slide 1
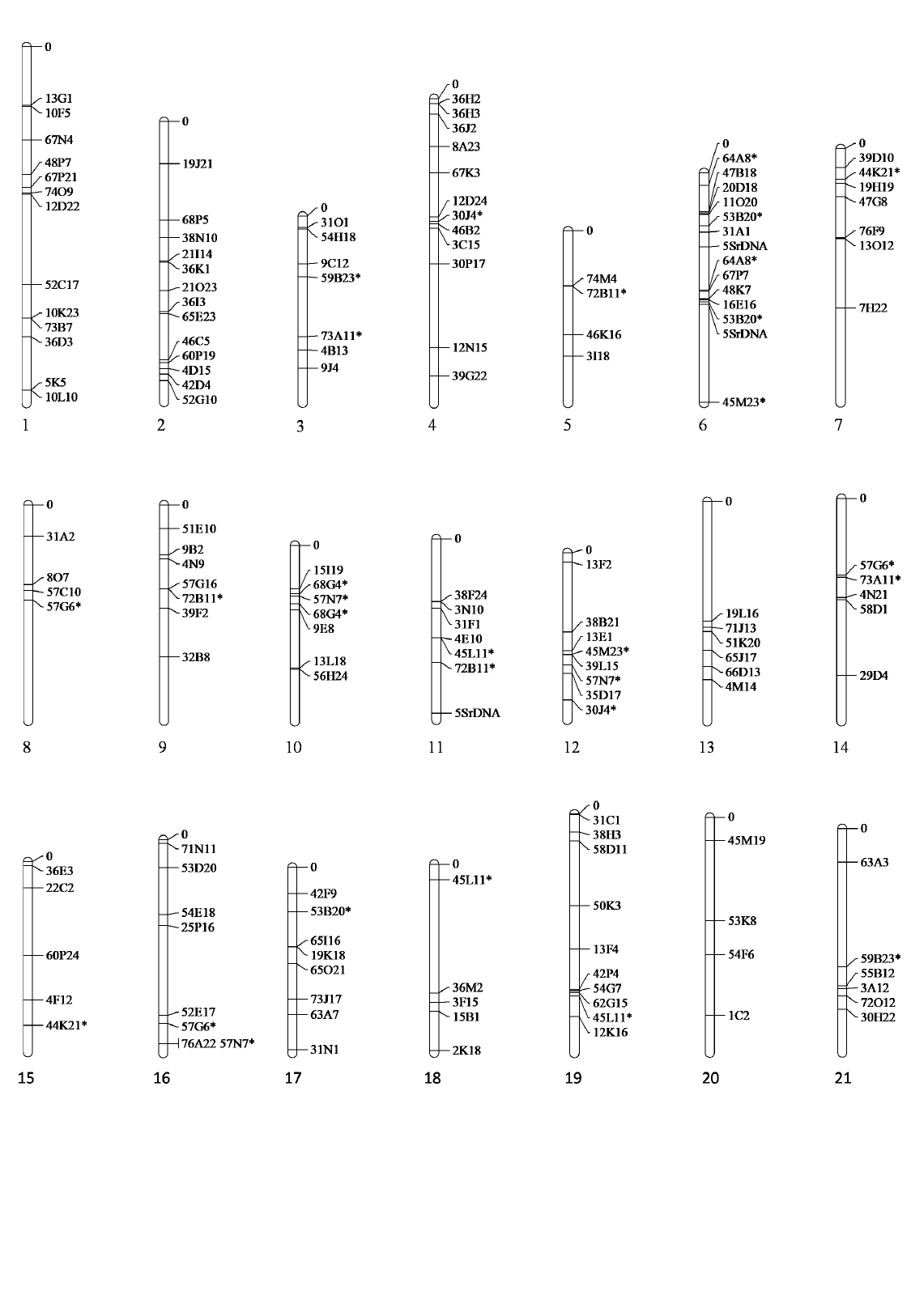

### Fig. S10

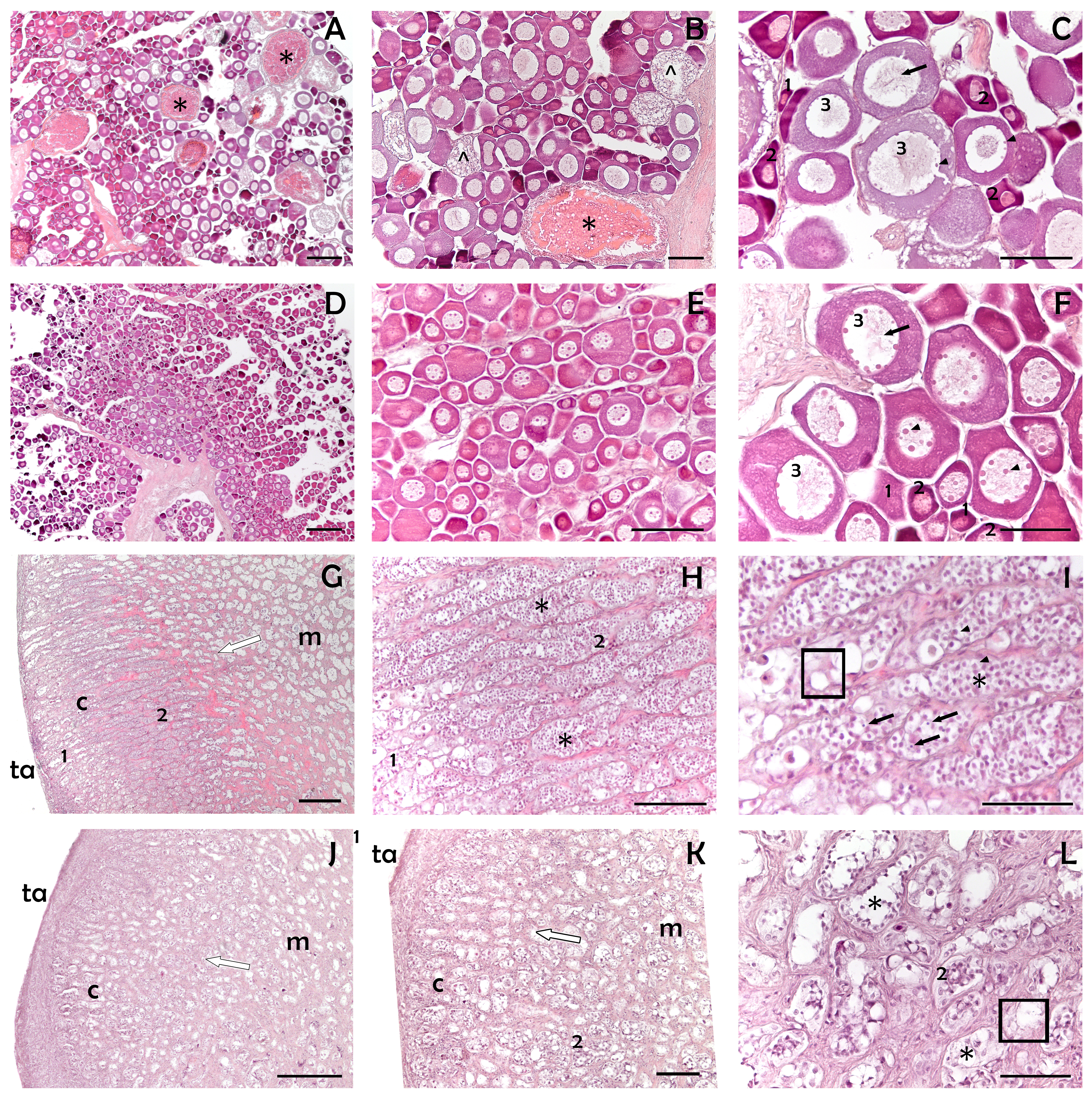

### Fig. S11

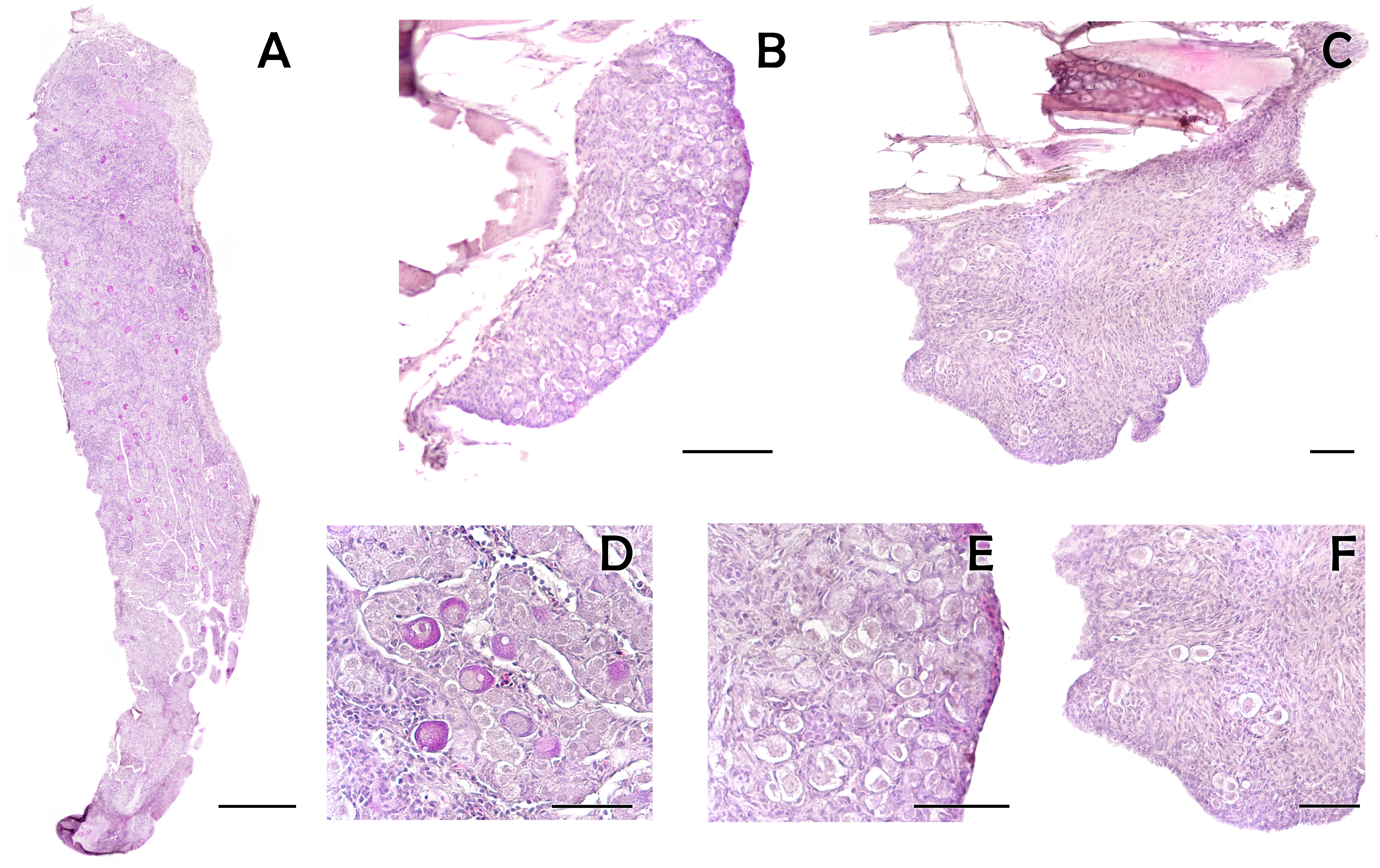

### Fig. S12

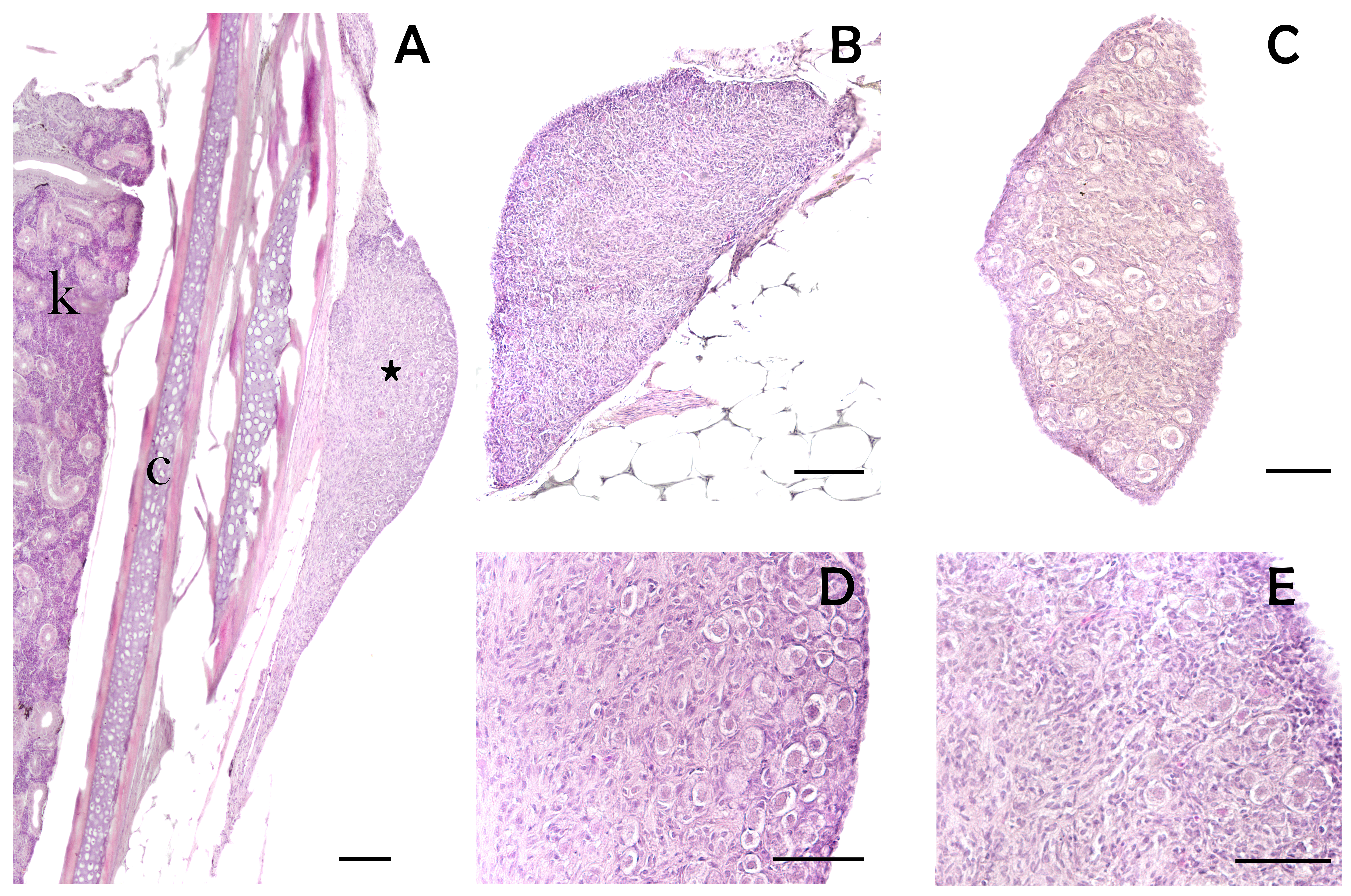

### Fig. S13

#### Slide 1
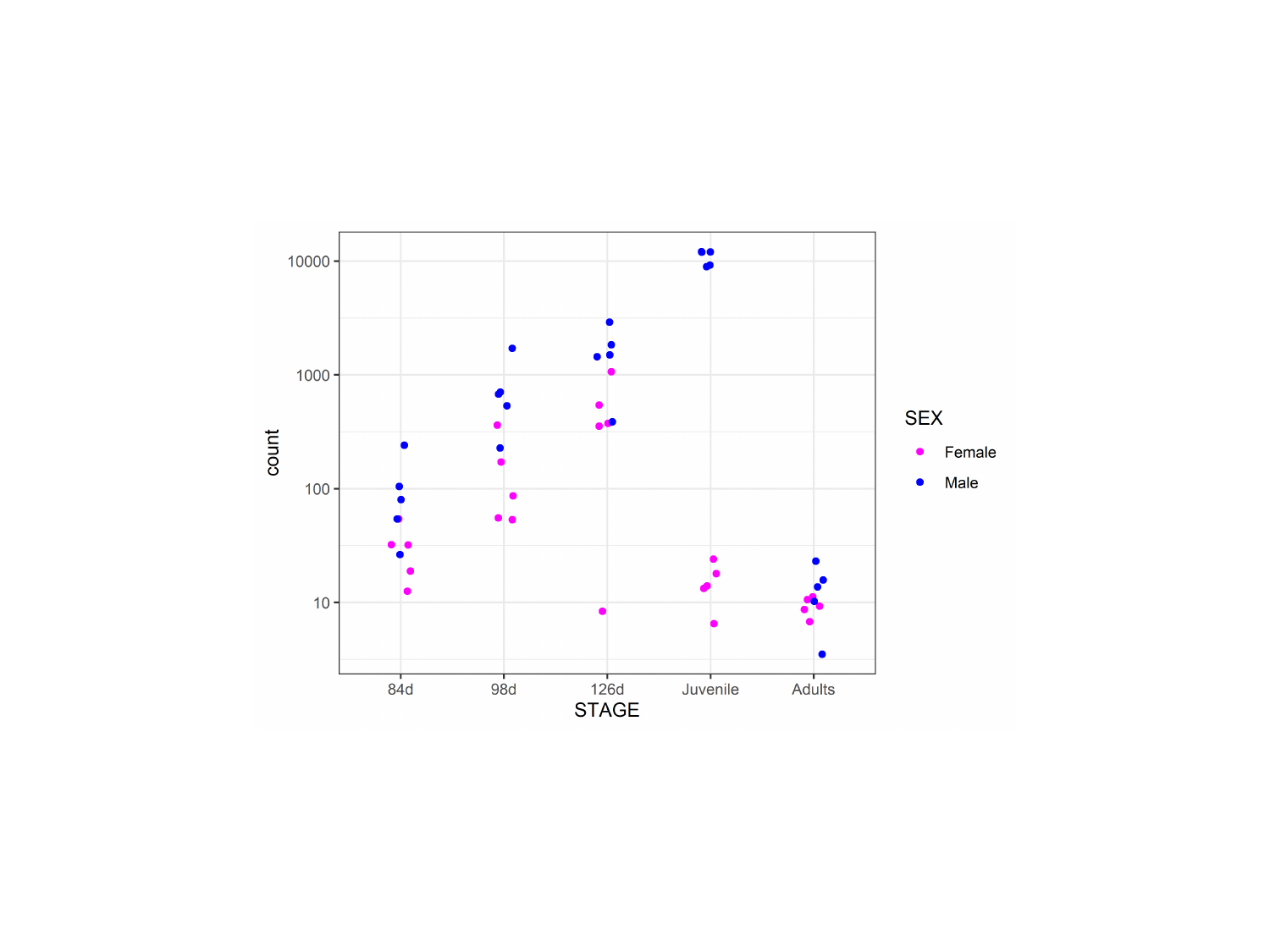

### Fig. S14

#### Slide 1
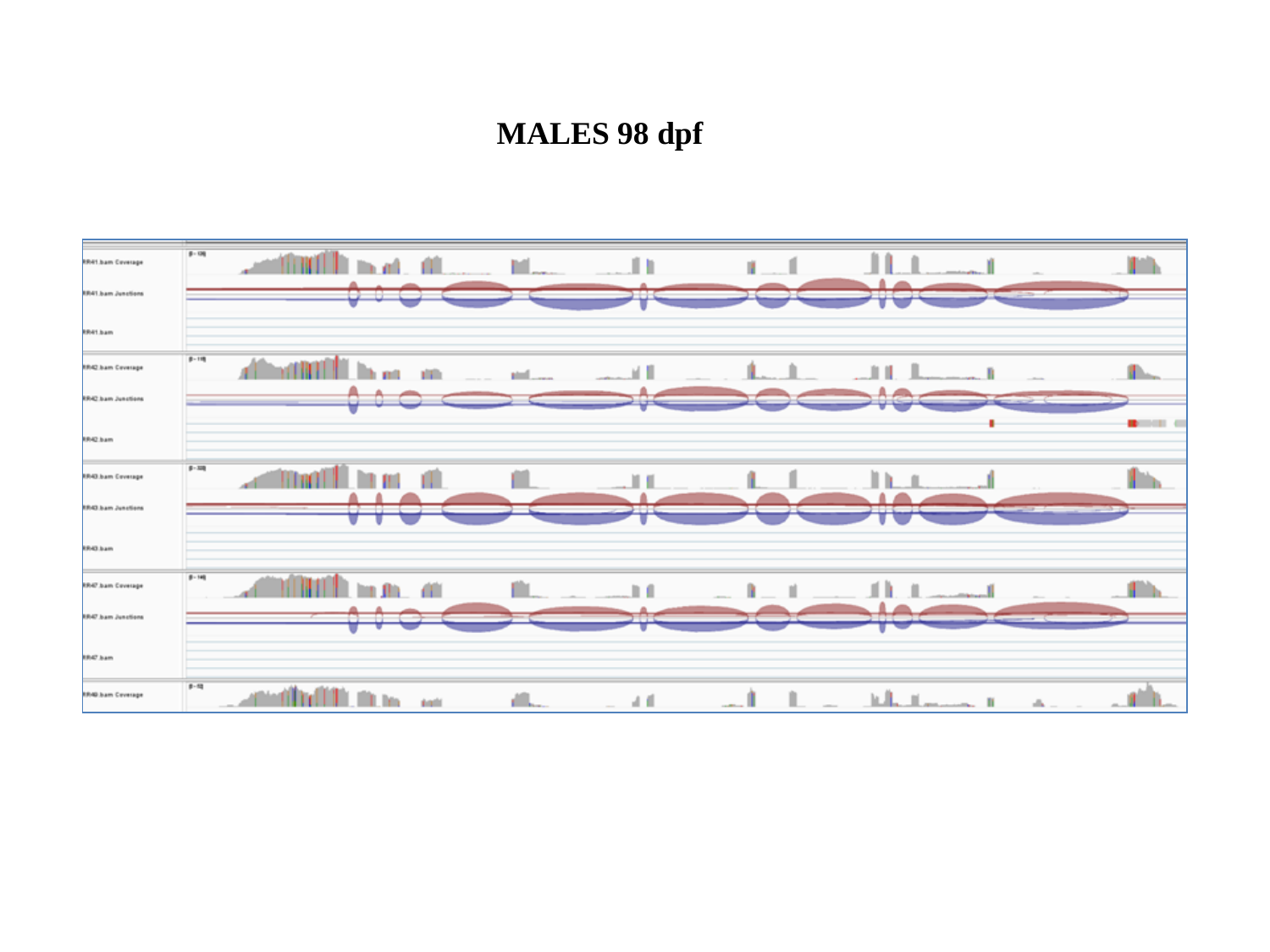

MALES 98 dpf

#### Slide 2
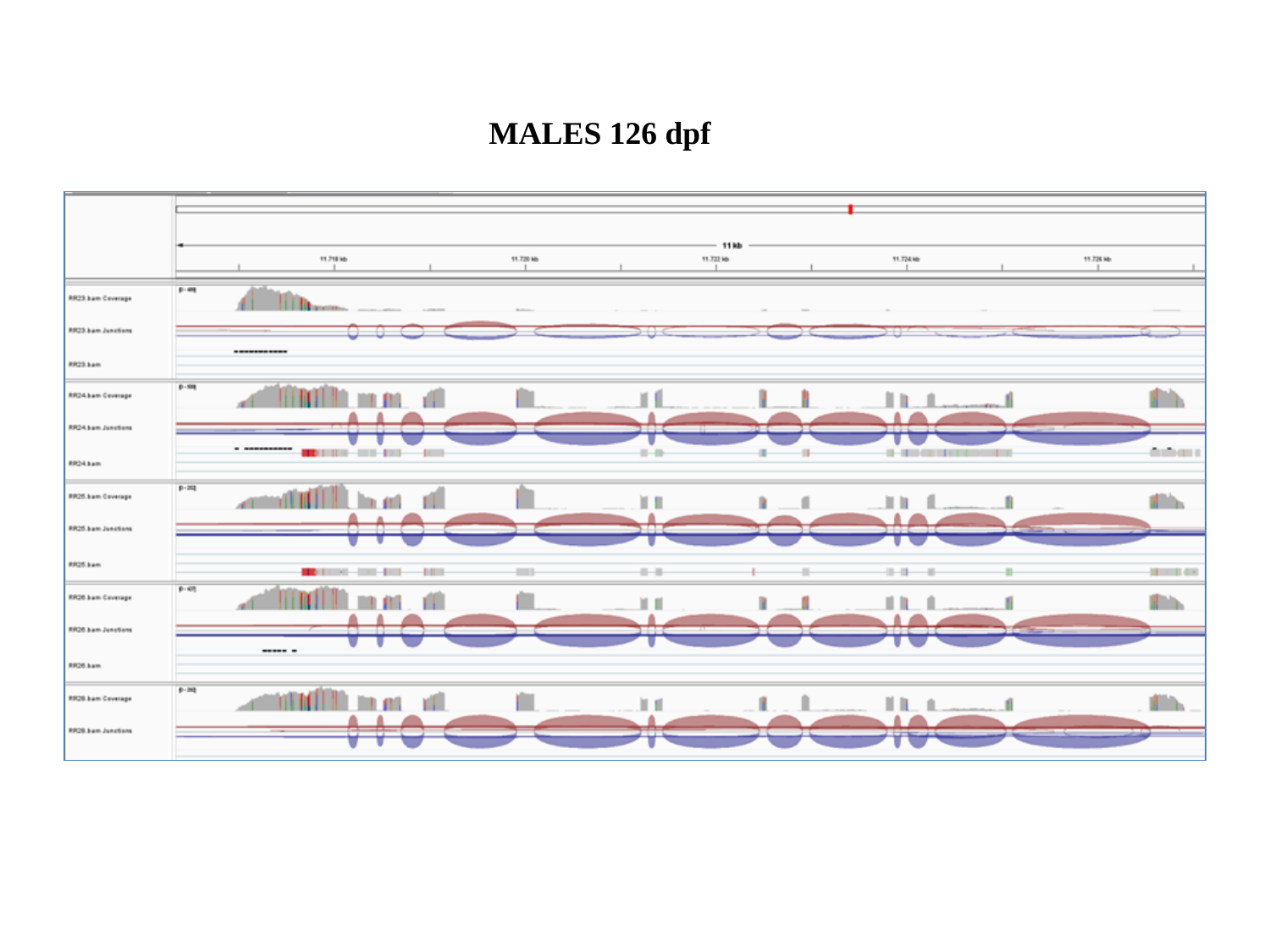

MALES 126 dpf

#### Slide 3
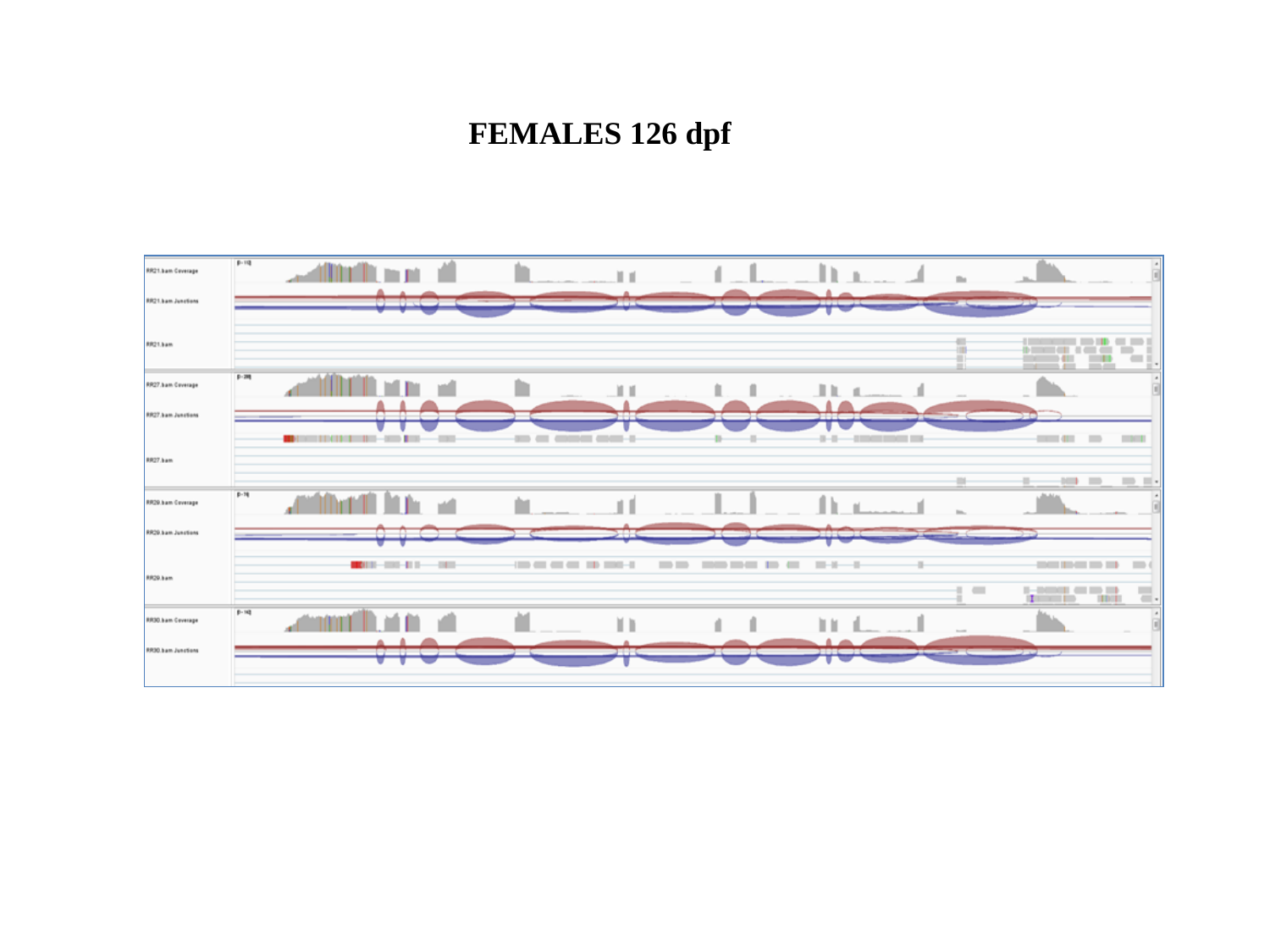

FEMALES 126 dpf

#### Slide 4
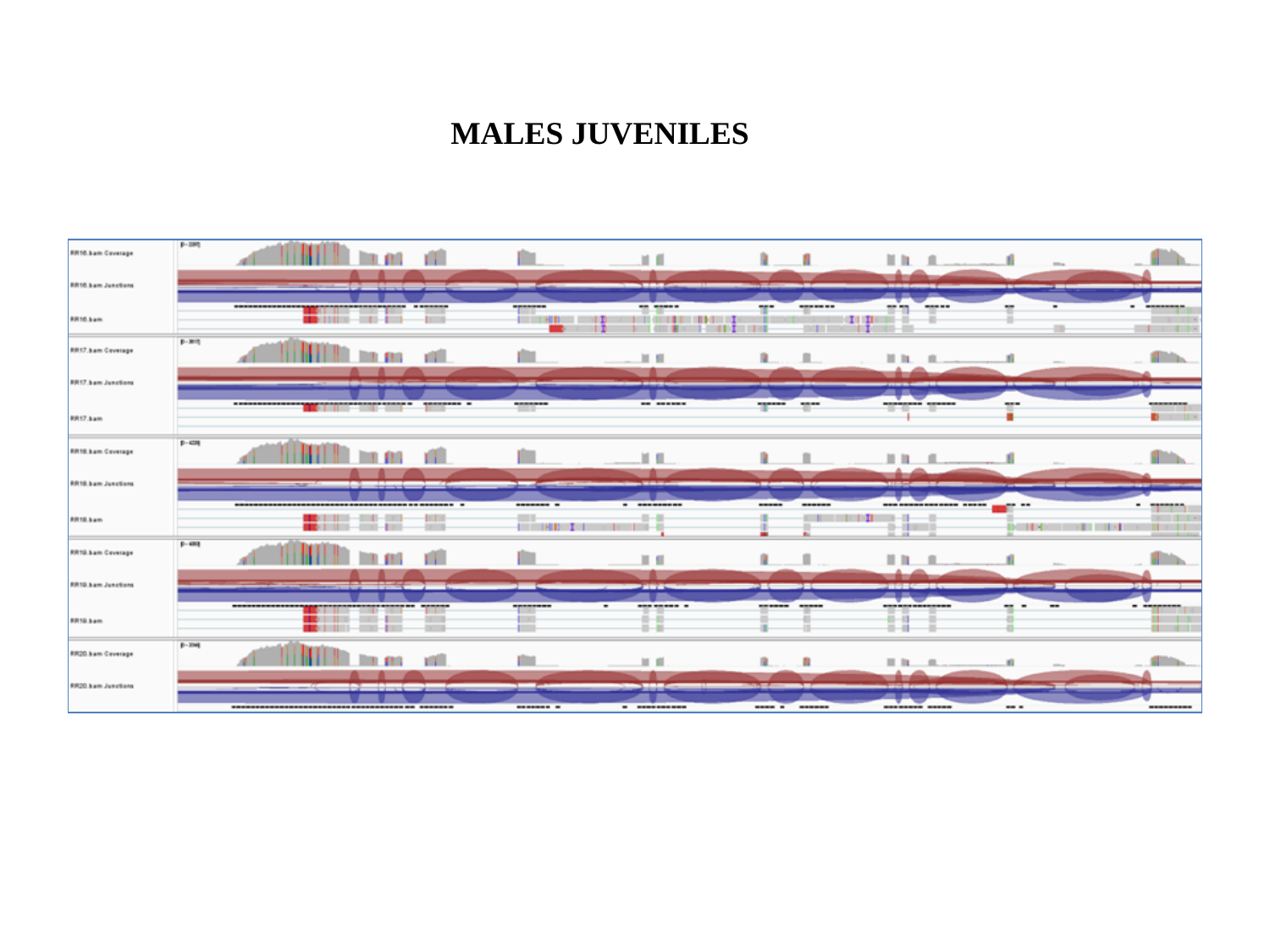

MALES JUVENILES

### Fig.S1

#### Slide 1
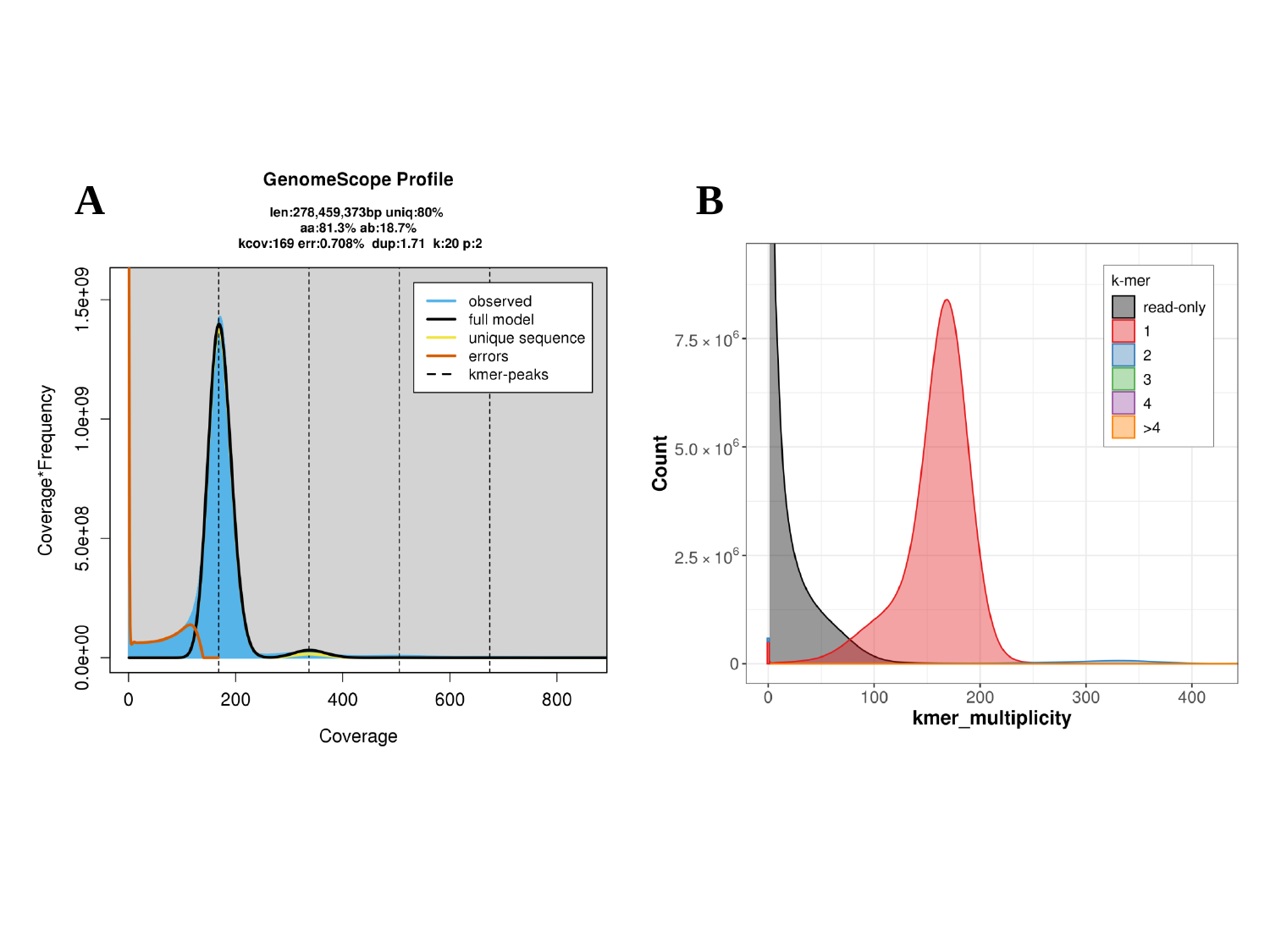

A
B

### Fig.S3A

#### Slide 1
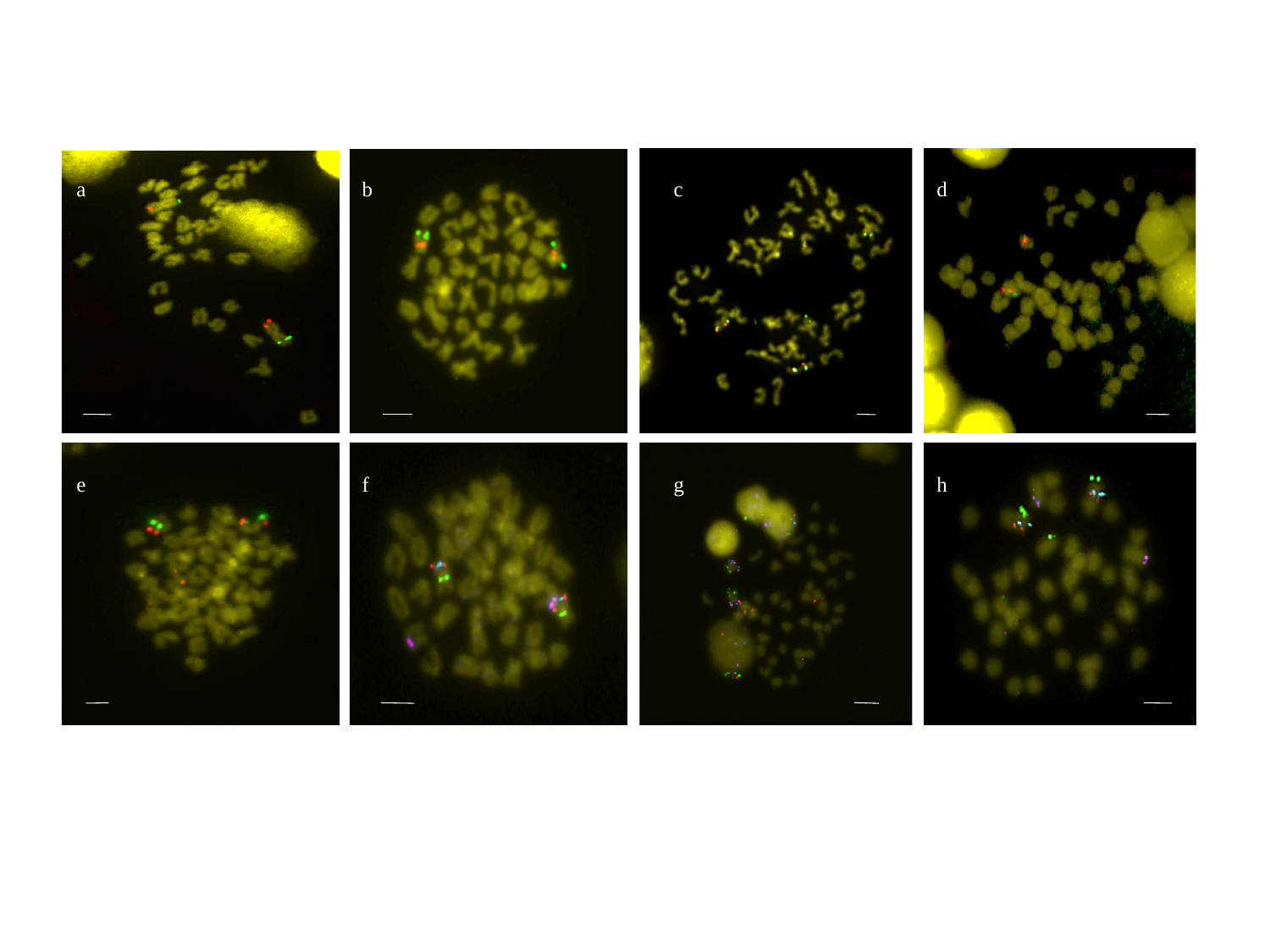

a
b
c
d
e
f
g
h

### Fig.S5

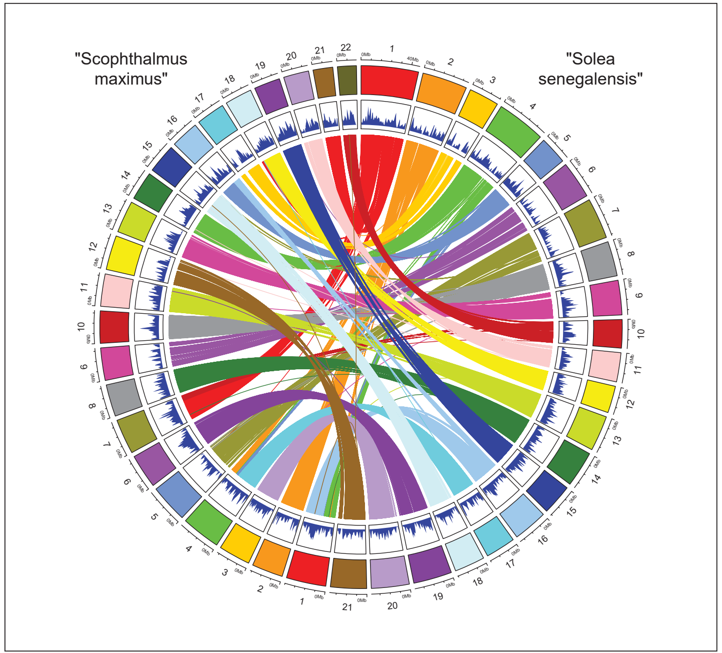

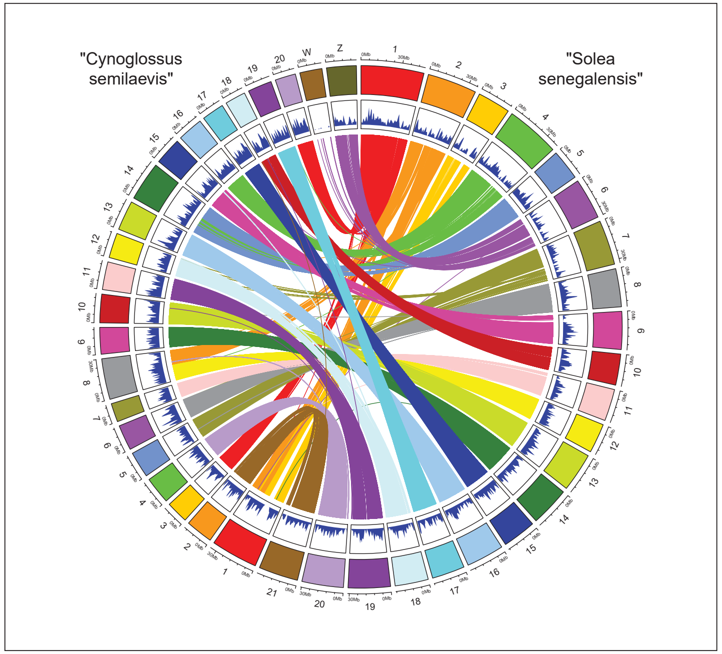

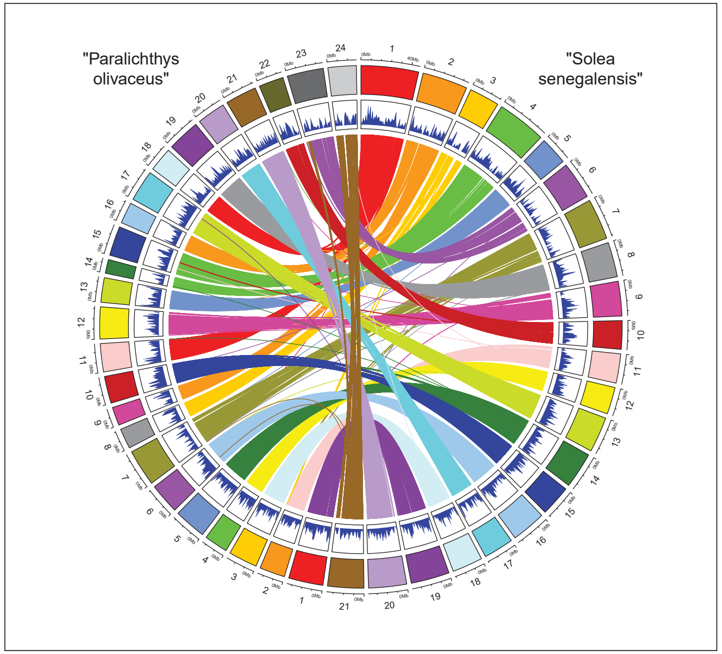

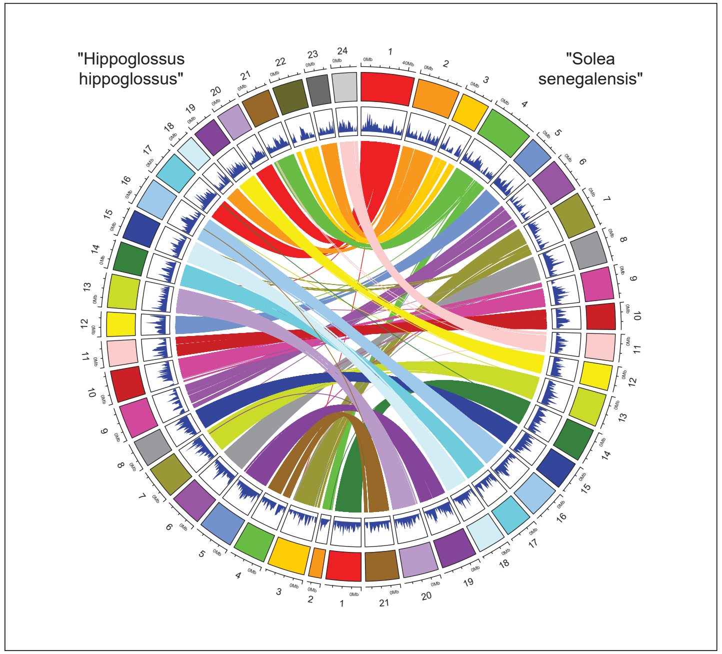

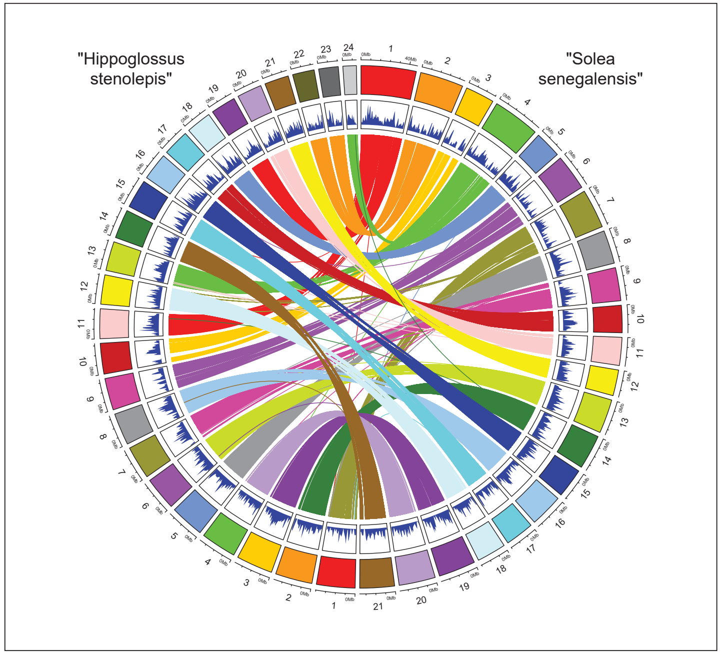

### Fig.S9

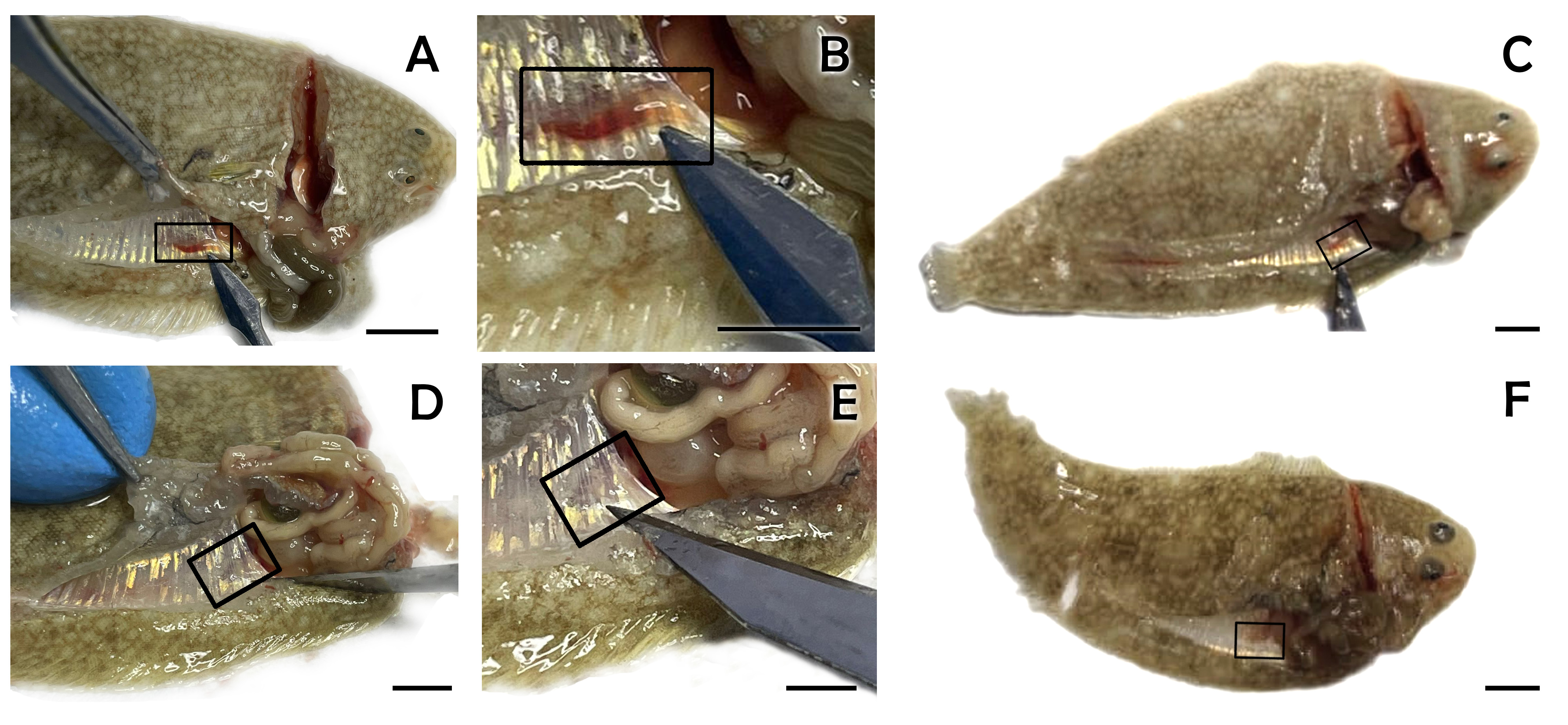

### Figs. 6, 7, 8

#### Slide 1
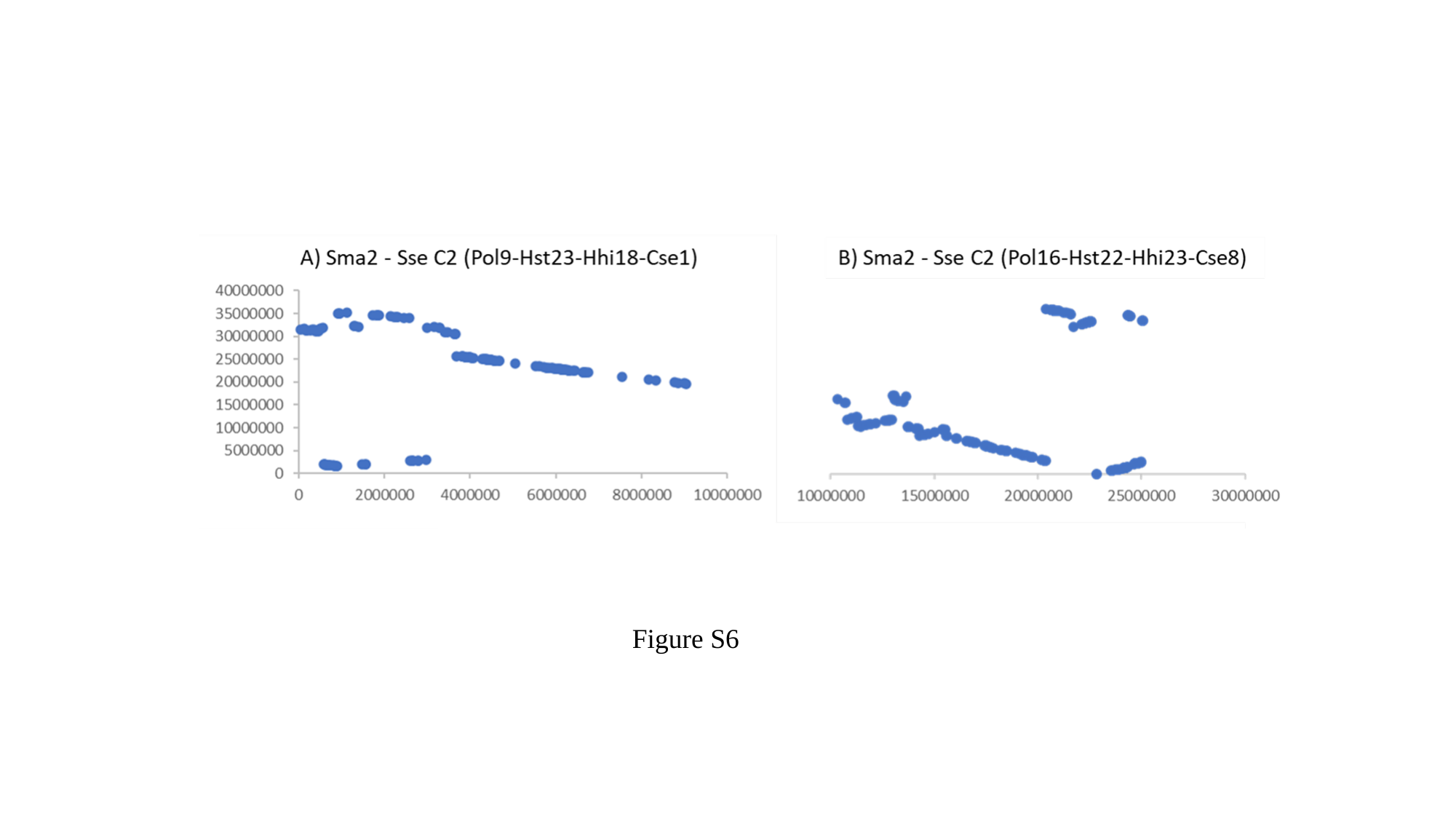

Figure S6

#### Slide 2
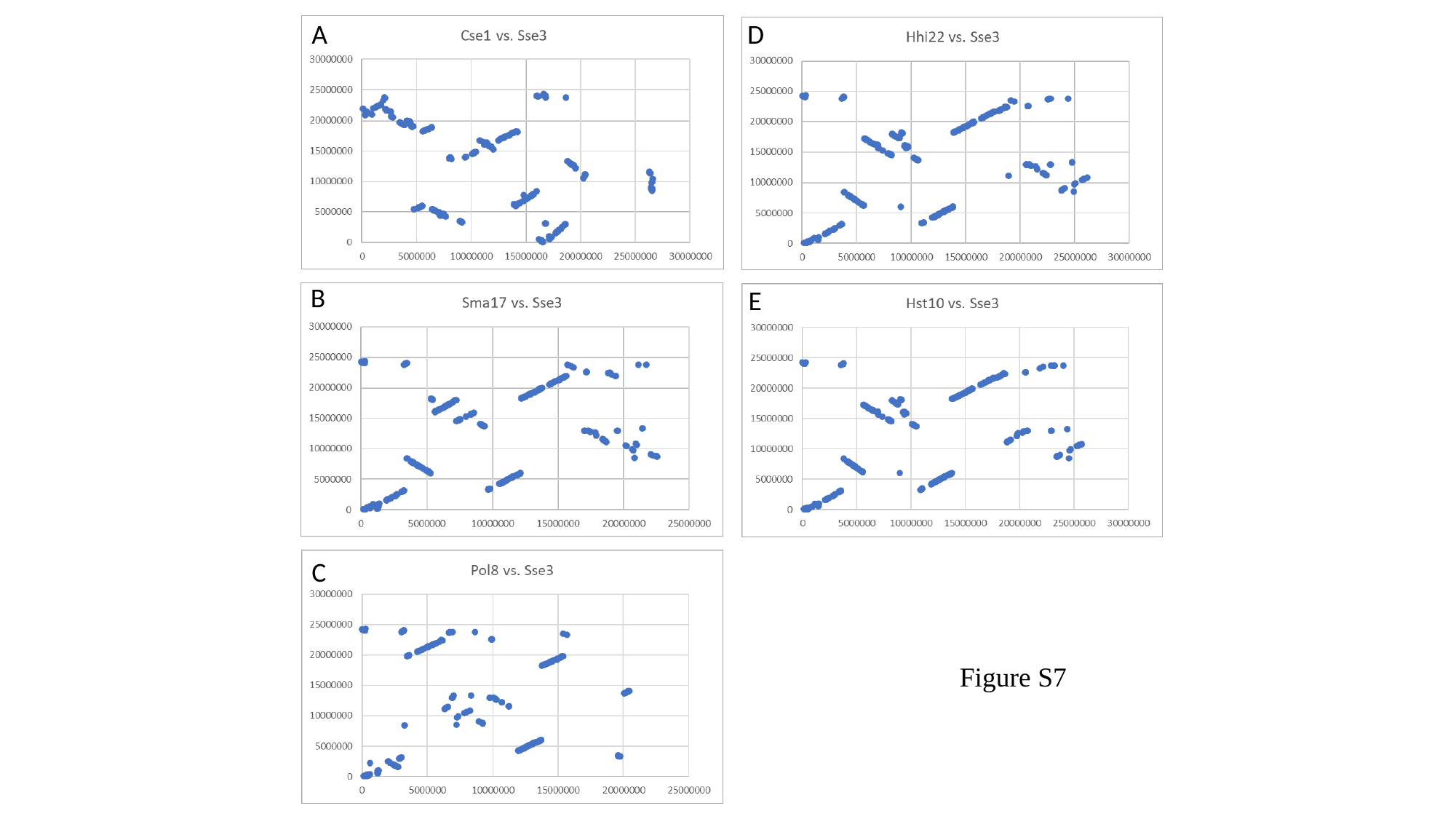

A
D
B
E
C
Figure S7

#### Slide 3
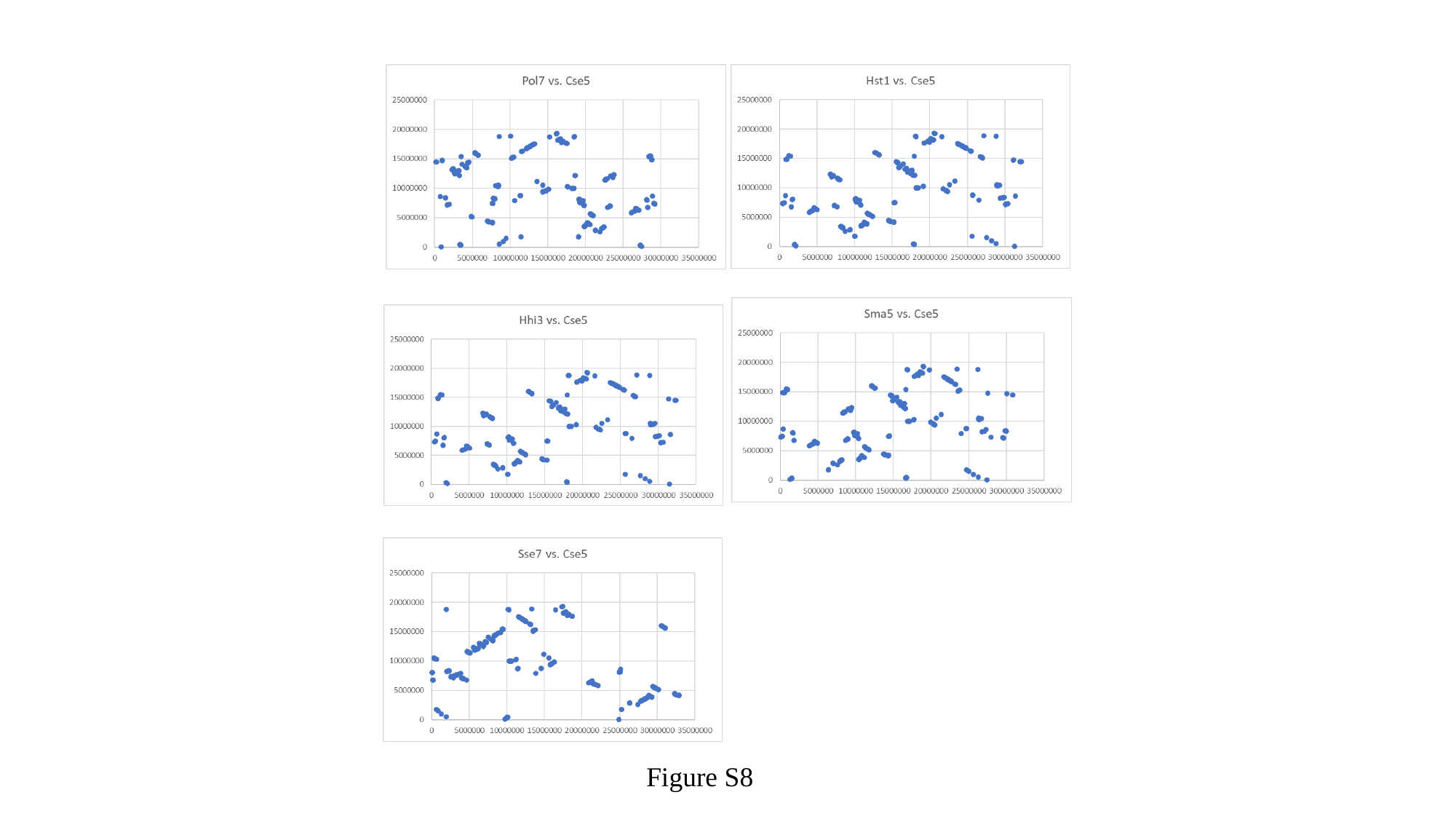

Figure S8
