## Supplementary material for "A chromosome-level genome assembly enables the identification of the follicle stimulating hormone receptor as the master sex determining gene in *Solea senegalensis*": Fig.S4

### Slide 1
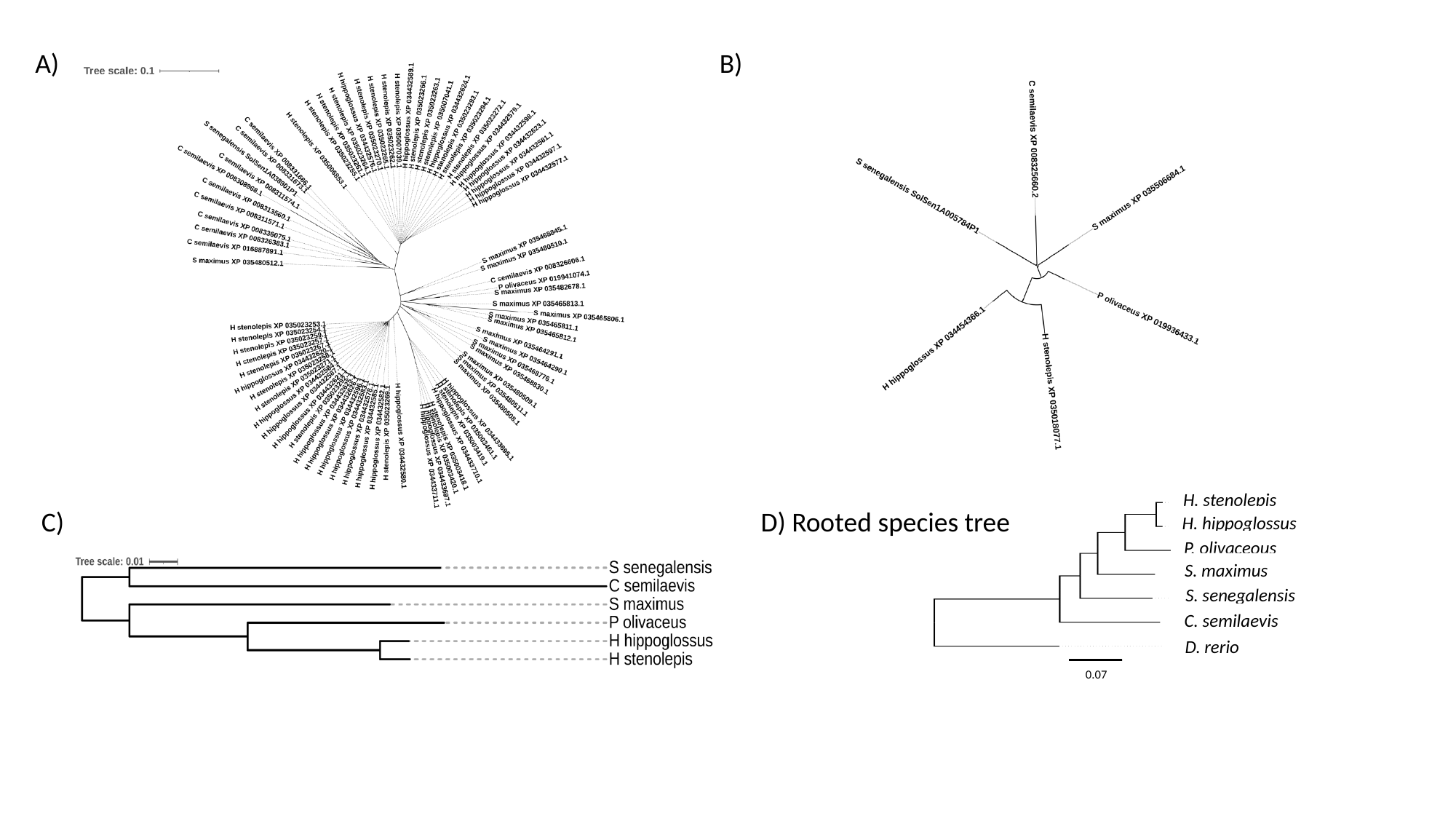

B)
 C) D) Rooted species tree
H. stenolepis
H. hippoglossus
P. olivaceous
S. maximus
S. senegalensis
C. semilaevis
D. rerio
0.07

### Slide 2
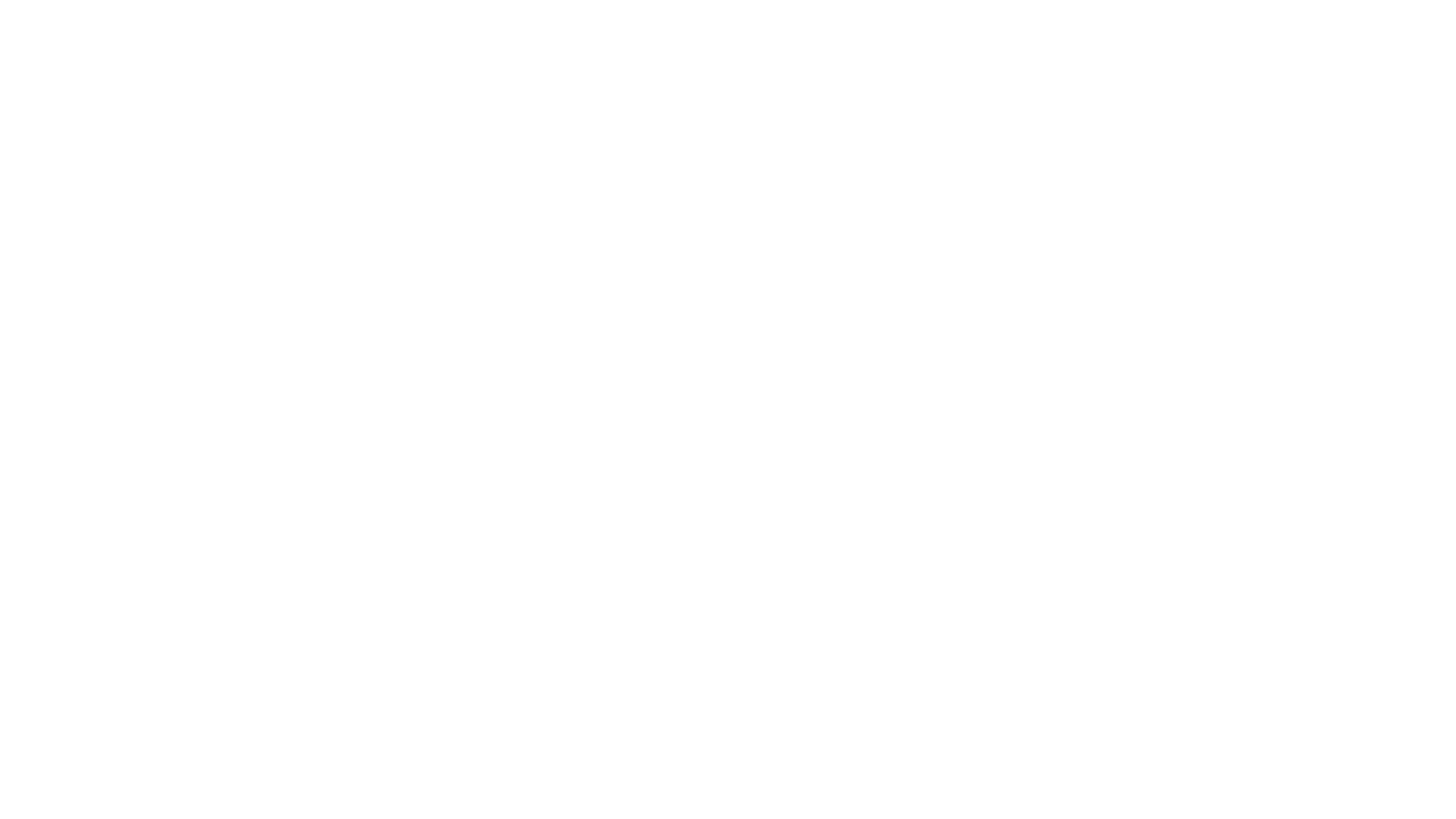
